# Proximity labelling reveals the compartmental proteome of murine sensory neurons

**DOI:** 10.1101/2025.06.13.659475

**Authors:** Julia R Sondermann, Allison M Barry, Feng Xian, Thomas Haberl, David Gomez-Varela, Fritz Benseler, Dietmar Schreiner, Nils Brose, Noa Lipstein, Manuela Schmidt

## Abstract

**Introduction:** Understanding the molecular architecture of peripheral sensory neurons is critical as we pursue novel drug targets against pain and neuropathy. Sensory neurons in the dorsal root ganglion (DRG) show extensive compartmentalization, thus understanding each compartment – from the peripheral to central terminals – is key to this effort.

**Methods:** To systematically profile this spatial complexity, we generated a TurboID^fl/fl^ transgenic mouse line (ROSA26^em1(TurboID)Bros^), enabling targeted proximity labelling and deep proteomic profiling of DRG neuron compartments via Tg(Advillin-Cre)^+^.

**Results:** Our data reveal distinct proteomic signatures across neuronal compartments that reflect specialized neuronal functions. We provide proteomic insights into previously inaccessible nerve terminals both in the periphery (innervating the skin) and in the spinal cord. Further, using a DRG explant model of chemotherapy-induced peripheral neuropathy, we uncover novel and discrete proteome changes, highlighting neuronal vulnerability.

**Conclusion:** Together, our findings provide a unique proteome atlas of the sensory neuron proteome across anatomical domains and demonstrate the utility of proximity labelling proteomics for detecting compartment-specific molecular alterations in a disease model.

## Introduction

Dorsal root ganglia (DRG) serve as critical hubs in the peripheral nervous system (PNS), housing the somata of sensory neurons that transmit information from the periphery (Middleton et al., 2021). DRG comprise sensory neurons as well as other cells, including glial and immune cells. Consequently, the interpretation of bulk proteomics data from DRG is inherently complex.

Adding to this complexity, DRG neurons exhibit spatial compartmentalization via axons to peripheral and central nerve terminals. Understanding their spatial proteome is therefore critical for elucidating the molecular mechanisms underlying sensory processing, neuronal maintenance, and pathological conditions such as chronic pain.

However, obtaining a comprehensive and spatially-resolved molecular profile of DRG neurons remains a significant technical challenge. Methods like fluorescence-activated cell sorting (FACS) can isolate specific cell types and have been used for synaptosome isolation (Biesemann et al., 2014), but FACS predominately prioritizes somata and/or nuclei. Isolating intact spatial compartments, particularly the nerve terminals, remains a major hurdle.

In recent years, proximity-dependent proteomic techniques like TurboID and APEX, have emerged as powerful tools for mapping subcellular proteomes and interactomes (Gingras et al., 2019; Qin et al., 2021). TurboID is an engineered, promiscuous biotin ligase that enables rapid and efficient labelling of proteins within a close proximity in living cells (Branon et al., 2018). To date, most applications of TurboID and its predecessor BioID have involved fusion proteins with compartment-specific targeting sequences, enabling selective biotinylation of proximal interactomes for downstream affinity purification and identification via mass spectrometry (MS) (Dingar et al., 2015; May et al., 2020; Roux et al., 2012). Unlike TurboID fusion proteins, the use of soluble TurboID variants enables unbiased labelling of the entire accessible proteome within a cell. Sunna et al. demonstrated that TurboID-mediated biotinylation covers approximately 59–65% of the proteome across cell lines expressing a cytosolic TurboID variant (Sunna et al., 2023).

Proximity labelling has also been successfully used to probe central nervous system compartments. For example, Rayaprolu et al. developed a mouse model to explore brain region-specific proteomic differences using TurboID *in vivo* (Rayaprolu et al., 2022). In another study, Hobson et al. employed APEX2 proximity labelling to map proteomic differences between somatodendritic and axonal compartments in acute brain slices, demonstrating correlates with respective neuronal function (Hobson et al., 2022).

Here, we generated a conditional mouse line expressing TurboID from the Rosa26 locus, enabling cell-type-specific proximity labelling *in vivo* using a Cre-lox system.

Paired to a Tg(Advillin-Cre)^+^ driver line (Zurborg et al., 2011), TurboID expression is prominent in DRG sensory neurons. Consequently, we were able to systematically profile the proteomes of DRG neuron compartments, providing a spatially resolved proteomic atlas. We then harnessed this technique to explore proteome changes after oxaliplatin treatment in an *in vitro* DRG explant model of chemotherapy-induced peripheral neuropathy (CIPN). CIPN is a common side effect of chemotherapy, with agents damaging sensory neurons and leading to painful neuropathy often preventing adequate cancer therapy (Sisignano et al., 2014). Indeed, we observed aberrant neuronal proteome dysregulation after oxaliplatin treatment, including changes not detected with bulk proteomics.

Together, our findings reveal novel insights into the compartment-specific protein distributions underlying key aspects of sensory neuron function and disease-related alterations.

## Methodology

### Generation of TurboID mice (ROSA26^em1(TurboID)Bros^) using CRISPR/Cas9

The conditional TurboID knock-in (cKI) mouse line was generated by site-directed CRISPR/Cas9 mutagenesis in the ROSA26 locus. Superovulated C57Bl6/J females were mated with C57Bl6/J males and fertilized eggs collected. In-house produced CRISPR reagents (hCas9_mRNA, sgRNAs, preformed Cas9_sgRNA-RNP complexes and the homology-directed repair fragment (HDR fragment)) were microinjected into the pronucleus and the cytoplasm of pronuclear-stage zygotes using an Eppendorf Femtojet (Eppendorf SE, Germany). Importantly, all nucleotide-based CRISPR/Cas9 reagents (sgRNAs and hCAS9_mRNA) were used as RNA molecules and were not plasmid-encoded, reducing the probability of off-target effects due to the short lifetime of RNA-based reagents (Doench et al., 2016; Tycko et al., 2019). The TurboID-containing HDR fragment was generated by subcloning a V5-tag-TurboID sequence into the Ai14-ROSA26-based plasmid backbone (Madisen et al., 2010) and isolated as a linear DNA fragment by SacI/AgeI restriction enzyme double digestion. Two sgRNAs targeting the ROSA26 Locus were selected using the guide RNA selection tool CRISPOR (Concordet & Haeussler, 2018; Haeussler et al., 2016). The correct locus-specific insertion of the HDR fragment was confirmed by localization PCRs with primers upstream and downstream of the HDR sequence. Extended details, including full sequences and genotyping strategy, are available in the Supplemental Methods.

### Animal husbandry, breeding, and biotin supplementation

Adult male and female TurboID^fl/fl^ control and Adv^Cre^;TurboID^fl/fl^ mice (12.5-26 weeks old for compartmental proteome, 57-71 weeks old for oxaliplatin part) and in-house bred wild type C57Bl6/J male mice (60-66 weeks old) were kept in a 12/12h day/night cycle with *ad libitum* access to food and water in an open-cage environment. The animal experiments were conducted in accordance with ethical guidelines and were approved by the Institutional Animal Care and Use Committee (IACUC) at the University of Vienna and the Austrian Ministry for Education, Science, and Research (BMBWF; license number 2021-0.138.925 amended by 2024-0.896.105). All procedures complied with the international ARRIVE guidelines and adhered to the European Union Directive 2010/63/EU on the protection of animals used for scientific purposes. No mice were removed from the study before their expected endpoint.

Breeding strategy: Male mice of the Tg(Advillin-Cre) driver line (referred to here as Adv^Cre^) (Barry et al., 2023; Lopes et al., 2017; Zheng et al., 2019; Zurborg et al., 2011) were crossed with here generated ROSA26^em1(TurboID)Bros^ (TurboID^fl/fl^) females to ultimately obtain mice heterozygous for Adv^Cre^ (Adv^Cre/+^) and homozygous for TurboID (referred to here as Adv^Cre^;TurboID^fl/fl^), along with their littermate controls (TurboID^fl/fl^). Full details are provided in the Supplemental Methods.

Biotin supplementation: TurboID^fl/fl^ control and Adv^Cre^;TurboID^fl/fl^ mice received biotin-enriched drinking water (0.22 mg/mL, prepared fresh every day) for 7 consecutive days to enhance biotinylation (Branon et al., 2018). On day 8, the mice were sacrificed. This dosing mirrored dosing of BioID mice, and can likely be reduced for TurboID, as reported for the C57Bl6/J -*Gt(ROSA)26Sor^tm1(birA)Srgj^*/J line (Rayaprolu et al., 2022).

Mouse numbers: The compartmental proteome was generated from pooled samples (3-6 animals per sample) from the following mice: TurboID^fl/fl^ control = 29M + 29F and Adv^Cre^;TurboID^fl/fl^ = 19M + 27F. Sample pools were stratified by sex.

Unilateral mouse DRG were used for explant cultures stimulated with Oxaliplatin, the DRG of the other side for the Vehicle condition in parallel. For the enrichment of biotinylated proteins in Adv^Cre^;TurboID^fl/fl^, 4 samples (3M, 1F) were prepared for each condition, with multiple mice pooled for each samples. In total, 10 males and 4 females were used per condition (3-4 mice per sample).

### Tissue dissection

Mice were sacrificed via CO_2_-inhalation and rapidly decapitated. The vertebral spine was removed for subsequent isolation of the dorsal part of the lumbar 3-5 spinal cord (SC) and lumbar 3-5 dorsal root ganglia (DRG). In parallel the sciatic nerves (SCN) of both sides were dissected. The plantar paw skin (dermis and epidermis, without muscle tissue) was removed by a 4 mm skin punch. Skin samples were immediately snap-frozen in liquid N_2_. All other tissues were washed in ice-cold 1× PBS to remove blood residuals before snap-freezing in liquid N_2_. All tissues remained at −80°C until further use.

### Protein extraction

All steps are done at room temperature (RT) except otherwise mentioned.

DRG: Tissue homogenisation was done in lysis buffer (LB; 2% SDS, 0.1 M tris, 5% glycerol, 0.01M DTT, 1× complete protease inhibitor cocktail) with 3x 10 strokes of a pestle/potter. To facilitate lysis the homogenate was sonicated using a Bioruptor Pico (10x 30 s ON/30 s OFF, 4°C, low frequency). All sonification steps mentioned below are done with the same settings (only cycle number varies). Samples were further incubated at 70°C, for 15’ min with 1500 rpm (Thermomixer C, Eppendorf). The remaining cell debris was pelleted by centrifugation (10’, 10000 ×g) and the supernatant was saved. All replicates of one condition were pooled and the concentration determined at 280 nm with a NanoPhotometer N60 (Implen, München, Deutschland).

SCN: Protocol was performed as above, except the centrifugation was followed by another 5 min-long centrifugation (RT, 10000×g) to further clear the lysate of lipid clouds. Before pooling and concentration measurement an acetone precipitation was done (see below).

SC: The DRG protocol was modified in that the pestle/potter step was omitted, instead the SC was cut into 3 smaller pieces and transferred to 2 mL LoBind tubes (prefilled with 500 µL of LB) for sonification Pico for 15 cycles. Other steps as for DRG, but before pooling and concentration measurement an acetone precipitation was done (see below).

Paw skin: the skin punches were cut into 4 smaller pieces, transferred into a potter prefilled with LB and homogenized with help of 3× 10 strokes of a pestle/potter. Lysis of tissue was facilitated by incubation for 20 min, at 70°C and 1500 rpm, followed by sonification for 15 cycles. The lysate was cleared of cell debris and excessive lipids by 2× centrifugation 10 min, 10000×g) and the supernatant was saved. Before pooling and concentration measurement an acetone precipitation was done (see below).

For the SCN, SC and paw skin an acetone precipitation was done to reduce the amount of lipids in the samples: 5× sample volume of pre-cooled (−20°C) 100% acetone was added to each sample and the tube was shortly inverted to ensure thorough mixing. If needed, samples were split into 2× 2 mL tubes to allow for 5× addition of sample volume. Precipitation of proteins was allowed for 3-3.5 h at −20°C. Next the precipitates were pelleted by centrifugation for 30 min and 14000×g. Pellets were washed with 1.5 mL pre-chilled (4°C) 80% ethanol and the centrifugation repeated as above. After removal of the ethanol, pellets were air-dried for 20-30 min. Proteins were reconstituted in LB by incubation for 15 min, 70°C, 1500 rpm. If were not fully solubilized, the incubation was repeated and LB volume increased until fully solubilized. Total protein concentration of pooled samples was determined with a NanoPhotometer at 280 nm.

### Streptavidin-mediated enrichment of biotinylated proteins

Streptavidin-affinity purification protocol is based on the high stringency protocol described in Go et al., 2021 with modifications. The required amount of streptavidin beads (Sera-Mag SpeedBead Blocked Streptavidin Particles, Cytiva, 83 µL bead slurry/ 1000 µg input) was transferred into LoBind protein tubes, washed twice with 500 µL LB without SDS, and resuspended in the same buffer. Protein lysates (input amount: DRG 1000 µg, SCN 600 µg, SC 800 µg, skin 2000 µg) were prepared to a final SDS concentration of 0.5%, final volume of 1500 µL. Lysates were incubated with the beads at 4°C for 3.5 h on an overhead shaker. Unless otherwise stated, all subsequent steps were performed at RT.

Following incubation, samples were placed on a magnetic rack for 2 min. Beads were then subjected to a series of washes, each performed with 1 mL of buffer, followed by overhead shaking for 1 min each (Grant-Bio PTR-35 Multi-Rotator) and incubation on the magnetic rack for 2 min until the solution cleared. The washes included one step with 2% SDS Wash Buffer (2% SDS, 25 mM Tris), two washes (three for skin samples) with modified RIPA buffer (50 mM Tris, 150 mM NaCl, 1.5 mM MgCl_2_, 1% Triton X-100, 1 mM EDTA), and three washes with 50 mM ammonium bicarbonate (ABC, pH 8). After the first ABC wash, the beads were transferred to a new 1.5 mL LoBind tube in 500 µL ABC buffer, followed by the addition of another 500 µL ABC before finalizing the washes.

For protein reduction, 135 µL of 5 mM dithiothreitol (DTT) in 50 mM ABC containing 0.8 mM biotin was added to each sample and incubated at 60°C for 30 min at 1000 rpm. Samples were then cooled to RT by briefly placing them on ice. Alkylation was performed by adding iodoacetamide (IAA) to a final concentration of 20 mM (15 µL of 200 mM IAA), followed by incubation for 30 min in the dark under constant agitation. The reaction was quenched by adding 1M DTT to a final concentration of 5 mM and incubating for 15 min with 3× 5 s vortexing steps in between. Proteins were digested for 17 h at 37°C and 950 rpm using 1.5 µg Trypsin/LysC per 1000 µg input. After the digest, peptides were eluted by placing the tubes on a magnetic rack for 2 min and transferring the supernatant to fresh 2 mL LoBind tubes. The beads were further washed with 20 µL MS-grade water (5 min, 500 rpm), and the wash was pooled with the initial eluate. Peptide concentrations were measured at 205 nm to calculate the required amount of SP3-beads (1:1 mix of Sera-Mag SpeedBead beads, 10-15 µL bead slurry to 40-50 µg peptides). To clean-up the peptides, peptide solution was split into two 2 mL tubes, the required bead amount and 100% ACN was added to reach 95% ACN. Samples were incubated for 8 min and 500 rpm, followed by a 2-min incubation on a magnetic rack. The ACN was discarded, and beads were rinsed with 400 µL 100% ACN, incubated for 2 min on a magnetic rack, and the ACN was discarded again. Peptides were reconstituted in 20-25 µL MS-grade water, incubated for 5 min with shaking at 500 rpm, followed by another 2-min incubation on a magnetic rack. The supernatants of previously split samples were pooled, centrifuged at 17000×g for 2 min, placed on a magnetic rack, and the final supernatant was transferred to a glass vial. Peptide concentrations were measured at 205 nm with a NanoPhotometer, and samples were acidified to a final concentration of 0.1% formic acid (FA). The samples were stored at −20°C until LC-MS measurement.

### DRG explants preparation

From both Adv^Cre^;TurboID^fl/fl^ and TurboID^fl/fl^ mice (fed with biotin-enriched water for 7 days, see above) 20 DRG per side were isolated and collected separately by side in two tubes containing DMEM/F-12 at RT for explant culturing (Fornaro et al., 2018). The DRG of one side were plated in 50 µL volume in to a well of a 24-well plate and incubated for 10 min at 37°C, 5% CO_2_. In the meantime, the oxaliplatin (100 µM) and/or vehicle (H_2_O, Veh) was mixed with DMEM +10% horse serum (HS) +100 ng/mL nerve growth factor (NGF) + 50 µM biotin and 500 µL/well were carefully added. The drug/Veh was incubated for 48 h at 37°C, 5% CO_2_.

To find an effective oxaliplatin concentration in line with previous reports (Livni et al., 2019), DRG explant cultures were prepared in a similar way from wild type mice. DRG from one mouse were separated into 4 wells for 4 conditions Veh, 25 µM, 50 µM and 100 µM oxaliplatin in DMEM+HS+NGF. The drug/Veh was incubated for 48 h at 37°C, 5% CO_2_.

### Sample preparation of DRG explants

The protein extraction was done similarly for both WT and TurboID DRG explants. All DRG of one well were transferred to an Eppendorf tube and washed twice with ice-cold 1× PBS by inverting the tube 10×. After discarding all PBS, 300 µL LB was added and DRG explants were sonicated for 10 cycles and further incubated at 70°C, for 15 min and 1500 rpm. Remaining cell debris was pelleted by centrifugation (5’, 10000×g) and the supernatant was saved. The total protein concentration was determined at 280nm with a NanoPhotometer N60. The TurboID samples underwent the above described streptavidin-affinity purification to enrich for biotinylated proteins and preparation of peptides (input 725 µg).

For the WT samples a modified version of the single-pot, solid-phase-enhanced sample preparation (SP3) method(Hughes et al., 2019) was employed to digest proteins and clean the peptides. In brief, 10 μL of pre-mixed Sera-Mag SpeedBead beads were added to 50 μg of protein. To facilitate protein binding to the beads, an equal volume of absolute ethanol was added, followed by incubation for 5 min at 24°C and 1000 rpm. Beads were collected on a magnetic rack for 2 min, the supernatant was discarded, and the beads were washed thrice with 500 μL of 80% ethanol. Subsequently, the beads were resuspended in 50 μL of 50 mM ABC, pH 8 containing 2 μg of Trypsin/Lys-C and incubated overnight at 37°C and at 950 rpm. Following protein digestion, ACN was added to each sample to reach a final concentration of 95%. After an 8-min incubation, beads were collected on a magnetic rack for 2 min. The supernatant was discarded, and the beads were washed with 900 μL of ACN. For peptide elution, the remaining ACN was allowed to evaporate for 2-3 min and the beads were resuspended in 40 μL of LC-MS grade water. Peptide concentrations were measured at 205nm using a NanoPhotometer N60. Finally, the peptide samples were acidified with FA to a concentration of 0.1% and stored at −20°C until LC-MS/MS analysis.

### Gel-electrophoresis & Western blot

All lysates from TurboID tissue that were going to be used as input samples for the streptavidin-affinity enrichment were checked by WB for the biotinylation success. The samples were prepared by dissolving them in 4× LDS Sample Buffer supplemented with 4× Reducing Agent to reach final 1×concentrations followed by heating to 70°C for 10 min. For protein separation, samples were loaded onto pre-cast NuPAGE® Bis-Tris 4– 12% gradient gels, the NuPAGE® (Life Technologies) electrophoresis chambers filled with 1× NuPAGE® MOPS buffer and the gels run for 40-50 min at 200 V.

Following electrophoresis, proteins were transferred onto PVDF membranes using the iBlot2® Gel Transfer Device (Life Technologies, program #0, 7 min). To block non-specific binding sites, membranes were incubated with 5% milk in PBS for 45 min at RT. Subsequently, membranes were incubated with primary antibodies (diluted in 1% milk/PBS; Stable 1) for 1.5 h, followed by an overnight incubation at 4°C with an additional 1:4 dilution. Membranes were washed with PBS (three short washes followed by three 5-min washes) before incubation with the respective secondary antibodies (diluted in 1% milk/PBS; see Stable 1) for 1 h. After washing with PBS as described before, immunolabeled proteins were detected using near-infrared fluorescence imaging (Bio-RAD ChemiDoc MP).

### LC MS acquisition

Nanoflow reversed-phase liquid chromatography (Nano-RPLC) was performed using NanoElute2 systems (Bruker Daltonik, Bremen, Germany) and coupled to a timsTOF HT mass spectrometer (Bruker Daltonik, Bremen, Germany) via a CaptiveSpray ion source. The mobile phase consisted of solvent A (100% water, 0.1% formic acid) and solvent B (100% acetonitrile, 0.1% formic acid).

500 ng peptides were loaded onto a C18 trap column (1 mm × 5 mm, ThermoFisher) before being separated on an Aurora™ ULTIMATE analytical column (25 cm × 75 µm) packed with 1.6 µm C18 particles (IonOpticks, Fitzroy, Australia) over a 60-minute gradient. The flow rate was maintained at 250 nL/min, except for the final 7 minutes, where it was increased to 400 nL/min. The gradient of mobile phase B was adjusted as follows: an initial linear increase from 2% to 15% over 30 minutes, followed by a rise to 35% over 22 minutes, and a sharp increase to 85% within 0.5 minutes. The flow rate was then adjusted to 400 nL/min in 0.5 minutes and held for 7 minutes to ensure complete elution of hydrophobic peptides.

Data acquisition was performed in data-independent acquisition (DIA) mode using parallel accumulation serial fragmentation (PASEF). Precursors with an m/z range of 350– 1100 were analysed across 12 MS/MS ramps in a cycle, with either two or three quadrupole switches per ramp. Each cycle contained 30 MS/MS windows within a 0.65–1.30 (1/k0) ion mobility range, using a fixed isolation window of 25 Da per step. The acquisition time per DIA-PASEF ramp was set to 100 ms, resulting in a total cycle time of approximately 1.38 seconds. Collision energy was ramped linearly from 65 eV at 1/k0 = 1.6 to 20 eV at 1/k0 = 0.6.

### Spectral deconvolution with DIA-NN

DIA-NN 1.8.2 was used to process DIA-PASEF data in library-free mode (Demichev et al., 2020). For the TurboID and WT samples, each tissue and, in case of Adv^Cre^;TurboID^fl/fl^ and TurboID^fl/fl^ genotypes were processed in a separate analysis step using the same spectral library. The spectral library was predicted from a Mus musculus proteome database from UniProt containing 17,070 protein entries (downloaded on 2021-07-08), in case of the TurboID samples it included in addition streptavidin (P22629|SAV_STRAV) and TurboID sequence. *In silico* digestion was performed using Trypsin/P, allowing up to two missed cleavages. A maximum of three variable modifications was allowed per peptide including methionine oxidation, and N-terminal acetylation, while Cysteine carbamidomethylation was set as a fixed modification. Peptide length for the search ranged from 5 to 52 amino acids. Precursor charge was set to 2-4. In accordance with the DIA-PASEF acquisition method, the m/z ranges were set to 350–1100 for precursor ions and 100–1700 for fragment ions. Mass accuracy was chosen automatically using the first run in the experiment. Protein inference was performed using unique genes. The single-pass mode was chosen. Quantification was carried out using RT-dependent cross-run normalization and the Quant UMS (high precision) option. The main DIA-NN output was processed using the R package DiaNN to extract MaxLFQ intensities (Cox et al., 2014; Demichev et al., 2020) for all identified protein groups, applying a q-value threshold of <0.01 at both the precursor and gene group levels. Matrices were merged by protein group were applicable for downstream analyses.

### Neuronal protein enrichments

Protein enrichments were calculated within each tissue/neuronal component prior to merging. First, proteins were filtered for distinct gene symbols, keeping the first instance where matches exist. Any protein detected in ≥75% of Adv^Cre^;TurboID^fl/fl^ samples for a given tissue, and no control sample was considered enriched (type “A”). This comprised the majority of enrichments. For proteins in ≥75% of Adv^Cre^;TurboID^fl/fl^ samples from any tissue, and any amount of control underwent differential expression testing using a standard limma framework (‘lmFit’ + ‘eBayes’) (Ritchie et al., 2015). Proteins were modelling by a grouped factor for Tissue and TurboID Status, and contrasted within tissue between TurboID and Control samples (ie. Paw.Turbo – Paw.Control). Proteins were considered enriched when FDR < 0.05, and LFC > 1 (“DEP”).

For explant analyses, we used the cross-tissue enrichments described above as a reference for neuronal enrichment, instead of setting a 75% threshold (ie. explant data were first filtered for neuronal enrichments using this reference list). Any proteins detected in the turbo samples and not control (type “not in TurboID^fl/fl^”), as well as any proteins that then hit a threshold of FDR < 0.05, and LFC > 1 (type “DEP”) were considered enriched in explant neurons.

### Pathway analyses

GO enrichments were performed on enriched proteins and differentially expressed proteins using the ‘enrichGÒ function from the ClusterProfiler and org.Mm.eg.db libraries in R (Wu et al., 2021). Both cellular component (CC) and biological pathways (BP) were extracted, with an adjusted p-value cutoff of 0.01 (Benjamini-Hochberg corrected). To account for redundancy in the GO terms, outputs were slimmed using default settings:

‘clusterProfiler::simplify(ego, cutoff=0.7, by=“p.adjust“, select_fun=min, measure = ’Wang’)‘.

For pathway analyses of TurboID-enriched proteins, a background of all proteins detected across the four tissue wildtype whole cell lysates + proteins detected with TurboID samples – this represents the entire universe of proteins which could have been enriched in neurons using the TurboID approach. For pathway analyses of differentially expressed proteins, a background of all proteins tested within the comparison was used. For pathway analyses of co-expression modules (below), module genes were compared to the universe of neuronal-enriched proteins.

### Synaptosome comparisons

Published DIA-MS synaptosome data from the skin/paw (Rawat et al., 2025) and spinal cord (Casañas & Montesinos, 2022) were compared to peripheral and central terminal proteomes respectively. Overlapping proteins were subject to pathway analyses via ‘ClusterProfiler‘as described above, against “CC” (cellular component). Human gene names were mapped to mouse using ‘biomaRt’ in R. Where > 10 pathways were significant, terms were filtered for “synap|signal|signalling|signaling|junction|membrane” for plotting, will all significant hits available in STable 3.

### Subtype enrichment

Neuronal subtype enrichment was calculated using an Overrepresentation Analysis (ORA) on previously published custom-made gene lists from the mouse DRG transcriptomics (Barry et al., 2023; Zheng et al., 2019). Briefly, custom gene enrichment lists for DRG neuronal subtypes were calculated from Zheng et al. (2019) using DESeq2 (Love et al., 2014; Zheng et al., 2019). ORA was then performed on TurboID-enriched proteins from each tissue using ‘Clusterprofiler ::enricher‘(Wu et al., 2021). For each tissue, the background was defined as any protein detected in whole cell lysate or TurboID samples, and the custom enrichment lists were filtered for proteins in the pre-defined background prior to ORA. ‘Clusterprofiler ::enricher’ does not allow the definition of both a custom background universe and custom gene set list (instead using only proteins defined in the custom gene sets as a background). To overcome this, the background proteins were defined as an additional gene set, and the gene set size was capped at 2000 to exclude this set from a direct enrichment analysis while maintaining them in the background. An FDR < 0.05 was considered significantly enriched, with an FDR =1 set for plotting.

### Immune cell enrichment

The depletion of immune cells was determined by the “infiltration score” from the ‘Im-muCellAÌpackage (0.1.0), a GSEA-based approach to query immune signatures (Miao et al., 2022). Protein names were converted to human gene symbols using the biomaRt package, and the expression of each tissue/compartment was presented against a background of all proteins detected in bulk and/or turbo samples, mirroring the pathway analysis (above). Missing values were imputed as zeros, and ImmuCellAI was run with default settings. The “infiltration score” was extracted from the resulting abundance matrix for plotting via ggplot2.

### Sexual dimorphism

Male/Female differences were tested using a standard limma framework for the neuronal proteome of each tissue (Ritchie et al., 2015). Proteins were modelled using a grouped factor for Tissue and Sex and contrasted within Tissue as M – F. Proteins were considered enriched when FDR < 0.05, and abs(LFC > 1). Phenotype permutations (eg. randomizing M/F labelling) was also employed via limma through ‘camera()‘as a more conservative test of pathway dysregulation. Pathways were considered enriched for an FDR < 0.05.

### Weighted co-expression network analysis

A weighted protein co-expression network was built using the WGCNA package in R (version 1.72-5) (Langfelder & Horvath, 2008). Turbo samples were filtered for the entire universe of enriched proteins (ie., enriched in any component). Proteins expressed only in a single component and proteins with a variance of 0 were then removed, resulting in a matrix of 3142 proteins across 39 samples.

Soft power was established for a signed network using ‘pickSoftThreshold(type = “signed“)‘, with a model fit of 0.8. The scale-free topology fit index failed for reasonable powers (< 30), suggesting a strong correlation among large gene groups, thus violating the assumption of the scale-free topology approximation. As noted in the WGCNA package documentation, this is commonly seen with datasets containing multiple tissue types, and a lack of scale-free topology fit does not inherently invalidate the data(Langfelder & Horvath, 2008). Consequently, we selected the recommended soft power of 14 for signed networks with 30-40 samples.

Network adjacency was calculated with ‘adjacency(type = “signed“)’ and the topological overlap matrix was calculated with ‘TOMsimilarity(TOMtype = “signed“)‘, both using default settings. Modules were established through adaptive branch pruning ‘dynamic-Mods <-cutreeDynamic(dendro = geneTree, distM = dissTOM, deepSplit = 2, pamRespectsDendro = FALSE, minClusterSize = 30)‘. Module eigengenes were then extracted for downstream analyses.

### Terminal protein-protein interaction networks

Neuronally enriched proteins (the “neuronal proteome”) for each terminal compartment were used to generate large-scale protein-protein interaction networks in Cytoscape, via the stringApp (Doncheva et al., 2023; Shannon et al., 2003). For the peripheral terminals, a core cluster of 433 proteins were highly connected through high confidence (0.7) protein-protein interactions (Szklarczyk et al., 2023). For the central terminals, 2271 proteins formed the core cluster. For each compartment, these networks were then clustered further. Here, community clustering was performed on the resulting STRING network in Cytoscape, where the Girvan-Newman fast greedy algorithm was implemented through ClusterMaker2 via GLay (Su et al., 2010; Utriainen & Morris, 2023).

Functional attributes within each cluster where then calculated via the stringApp, against a background of the entire peripheral/central terminal proteome respectively. Cluster numbers are somewhat arbitrary but correspond to the supplemental table outputs with full enrichment details.

### Oxaliplatin effects

The effects of the chemotherapy agent oxaliplatin on explant cultures was tested using limma. For DRG explants dissected from wildtype mice, oxaliplatin was administered at 25 µM, 50 µM and 100 µM concentrations (or vehicle). Proteins here were modelled as Concentration.Condition, and contrasted within each concentration. For TurboID mice, data were first filtered for Turbo samples and TurboID-enriched proteins prior to modelling by Condition. In each case, limma was used to contrast by treatment condition (oxaliplatin – vehicle). Proteins were considered differentially expressed with FDR < 0.05, and abs(LFC) > 0.5. Downstream processing is described in the ‘*Pathway analyses*‘section.

Comparisons against LFC and overall expression were compared to previously published, DIA-MS from *in vivo* oxaliplatin dosing (Yang et al., 2022)

### Rank-Rank Hypergeometric Overlap

Rank-Rank Hypergeometric Overlap (RRHO) testing was performed on limma outputs. Here, concordant and discordant gene profiles were calculated via the Stratified method using ranked ‘-log_10_(p-values) * sign(effect size)‘, implemented through the RRHO2 library in R in line with their published recommendations (Cahill et al., 2018)

### STRING analysis

STRING protein-protein interaction analyses were performed through Cytoscape 3.10.3 using the stringApp (Doncheva et al., 2023; Shannon et al., 2003) A list of DEPs for each significant biological pathway were extracted from the ClusterProfiler output (see: *Pathway analyses*) and loaded into the stringApp with default settings. Proteins were coloured based on differential expression in either TurboID only, or TurboID + bulk explant datasets. Drug targets were determined via https://dgidb.org/, filtered for “regulatory approval == Approved” (Cannon et al., 2024).

### Immunohistochemistry (IHC)

For IHC the tissue was dissected as above with modifications: instead of the dorsal part of the SC, the whole L3-5 SC was taken out. The plantar paw punch contained not only the epidermis and dermis but all layers below the bones. The dissected tissue was fixed in 4% formalin/PBS for 3 h at 4°C, followed by overnight cryoprotection in 30% (w/v) sucrose at 4°C. For sectioning, the tissue was embedded in Tissue-Tek O.C.T. compound and frozen. Serial sections (SCN, DRG: 10 μm, SC: 30 μm, skin: 20-40 μm) were obtained using a cryostat and mounted on SuperFrost Plus slides. Sections were stored at −80°C until further processing.

Frozen sections were thawed at RT for 30 min before permeabilization and blocking in 0.4% Triton X-100/PBS containing 5% donkey serum for 30 min. Primary antibodies, diluted in 0.1% Triton X-100/PBS with 1% donkey serum, were applied overnight at 4°C. Following three washes in PBS, sections were incubated at RT for 2 h with Alexa Fluor-conjugated secondary antibodies in PBS containing 1% donkey serum and 0.1% Triton X-100. Unbound antibodies were removed by washing with PBS, and sections were mounted using SlowFade™ Glass Soft-set Antifade Mountant Medium containing 4′,6-Diamidino-2-phenylindole (DAPI). Antibodies used are listed in STable 1.

### Graphics

R base functions and the following libraries were used to compile plots unless otherwise described: ggplot2, ggbiplot, ComplexHeatmap, cowplot, gridExtra. Figures and schematics were designed in Inkscape.

### Use of generative AI

Generative AI tools (OpenAI ChatGPT) were used solely to improve phrasing, clarity, and formatting. All scientific concepts, experimental designs, hypotheses, and project goals were developed by the research team. No synthetic or AI-generated data were included.

## Results

We profiled the compartmental proteome of murine sensory neurons of DRG using the well characterized Tg(Advillin-Cre) driver line (referred to here as Adv^Cre^) (Barry et al., 2023; Lopes et al., 2017; Zheng et al., 2019; Zurborg et al., 2011) crossed with the here generated ROSA26^em1(TurboID)Bros^ (TurboID^fl/fl^) mouse line (Figure 1; please see Methods for details on line generation, breeding, and husbandry). In Adv^Cre+^;TurboID^fl/fl^ mice (referred to here as Adv^Cre^;TurboID^fl/fl^), TurboID is expressed in the cytoplasm of Advillin-positive cells, resulting in promiscuous biotinylation of intracellular proteins, largely within sensory neurons (Fig 1A). Respective control tissue was derived from littermates lacking Adv^Cre^ (Adv^Cre-^;TurboID^fl/fl^ , referred to as TurboID^fl/fl^). TurboID^fl/fl^ samples do not express TurboID and therefore serve as controls to account for endogenous biotinylation (Fig 1A). Tissue from wildtype mice were also collected for downstream comparisons (Fig 1B).

**Figure 1.**
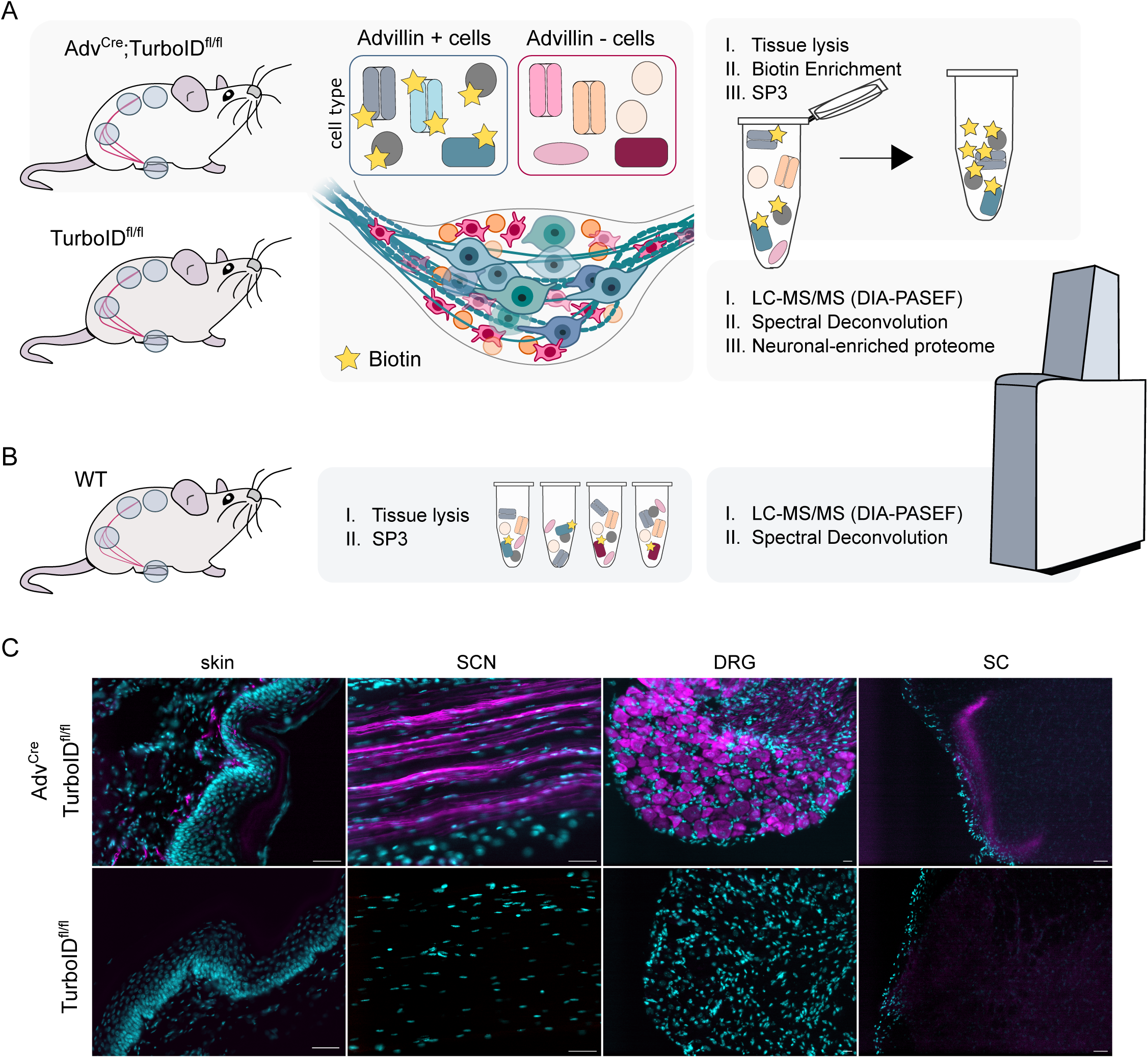
Adv^Cre^;TurboID^fl/fl^ mice were used to target the DRG neuron proteome. A. Schema overview for biotinylation of DRG proteins. Adv^Cre^;TurboID^fl/fl^ and TurboID^fl/fl^ mice were provided biotin-enriched water; in Adv^Cre^;TurboID^fl/fl^, this promotes the biotinylation of proteins within Advillin-expressing (+) cells. Advillin-negative (-) cells and tissues from TurboID^fl/fl^ mice only contain endogenously biotinylated proteins. Tissues were lysed and biotinylated proteins were purified followed by downstream LC-MS/MS on a timsTOF HT in DIA-PASEF mode. B. Bulk proteomes were isolated from wildtype (WT) mice for each corresponding tissue in A. C. Representative immunohistochemistry from TurboID^fl/fl^ (control) and Adv^Cre^;TurboID^fl/fl^ mice across tissues. Cyan, DAPI. Magenta, Streptavidin-555 (labelling biotin). Scale bar = 50 µm.

DRG neurons are pseudounipolar, with peripheral branching to target organs and central branching to the spinal cord. To cover this spatial complexity, we isolated tissue from the spinal cord (SC), dorsal root ganglia (DRG), sciatic nerve (SCN) and paw skin after providing biotin in drinking water for 7 days (Branon et al., 2018; Rayaprolu et al., 2022). Biotinylated proteins were enriched after tissue lysis followed by analysis using liquid chromatography-coupled mass spectrometry (LC-MS/MS). Biotinylation in each tissue of Adv^Cre^;TurboID^fl/fl^ mice was confirmed by immunohistochemistry (Fig 1C, SFig 1A) and validated via Western blotting (SFig 1B). As expected, we see a clear enrichment of biotin labelling (visualised via biotin-streptavidin) across all compartments (Fig 1C).

Using Western blotting, biotin signal was readily detected in tissue lysates but was less prominent in skin (SFig 1). The latter may be due to low protein quantity after lysis, while TurboID-positive afferents were evident by immunohistochemistry (Fig 1C, “skin”).

To generate a comprehensive and quantitative compartmental proteome atlas of DRG neurons, we employed data-independent acquisition combined with parallel accumulation–serial fragmentation (DIA-PASEF) mass spectrometry on Adv^Cre^;TurboID^fl/fl^ and Adv^Cre^ -negative control tissue (TurboID^fl/fl^) (Fig 2, SFig 2-5, STable 1). Both overlapping and distinct proteins were detected across DRG compartments (Fig 2A). A comparison of Adv^Cre^;TurboID^fl/fl^ with TurboID^fl/fl^ controls shows more protein (SFig 2A) and more protein groups (SFig 2B) in Adv^Cre^;TurboID^fl/fl^ samples. Samples also clustered by Adv^Cre^;TurboID^fl/fl^ compartment (Fig 2B) and across genotype (Fig 2C; SFig 2C). A full expression matrix is available in STable 1.

**Figure 2.**
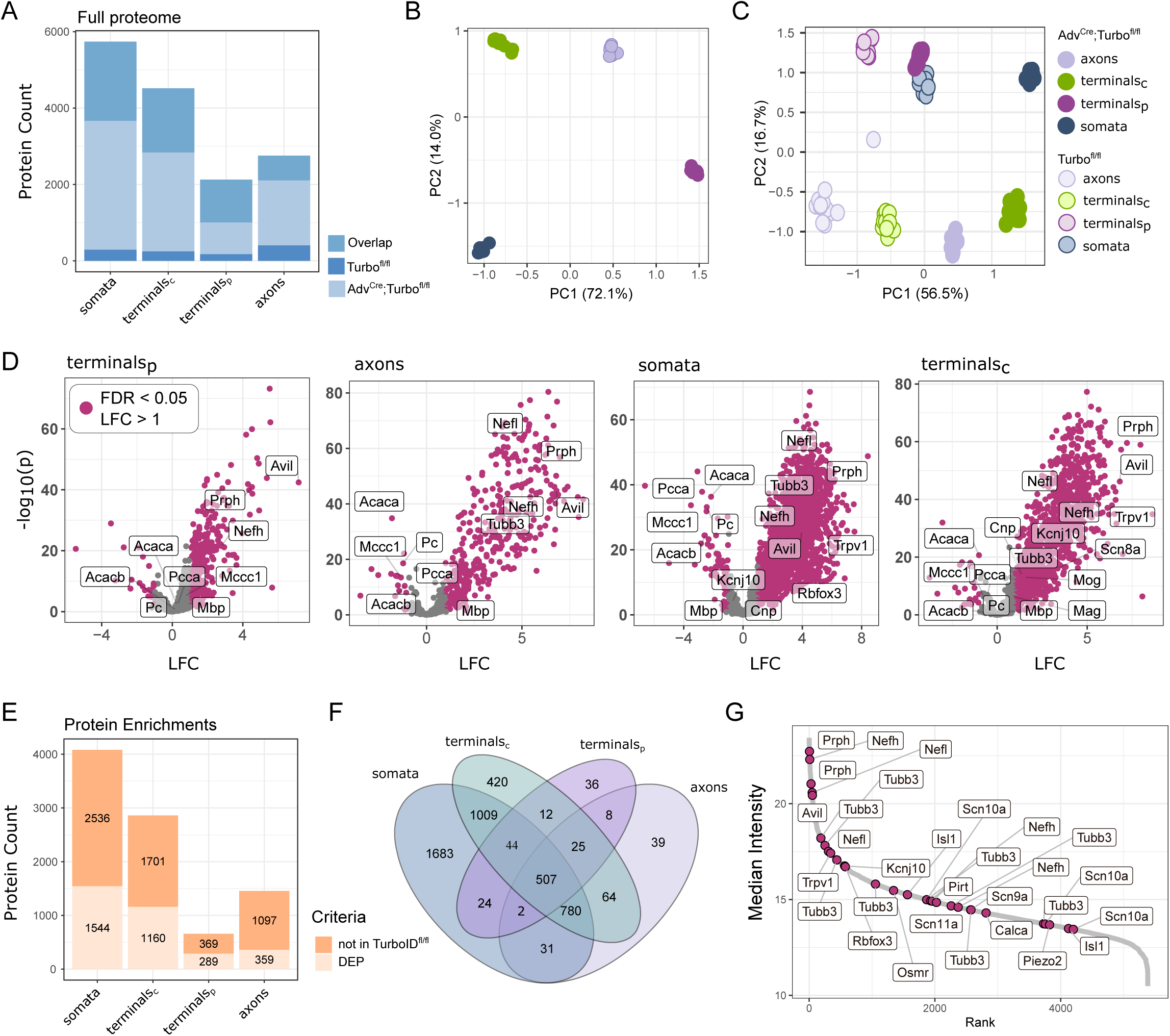
Protein enrichment by TurboID. A. Protein counts by sample, with “overlap” representing proteins detected in both Adv^Cre^;TurboID^fl/fl^ and TurboID^fl/fl^ control tissue. B-C. PCA for Adv^Cre^;TurboID^fl/fl^ (B) and both Adv^Cre^;TurboID^fl/fl^ and TurboID^fl/fl^ (C). D. Volcano plots highlight DEPs based on their LFC in Adv^Cre^;TurboID^fl/fl^ versus TurboID^fl/fl^. E. Numbers of enriched proteins, including both DEPs from D (Adv^Cre^;TurboID^fl/fl^ vs TurboID^fl/fl^), as well as proteins detected in Adv^Cre^;TurboID^fl/fl^ but not TurboID^fl/fl^ controls (“not in TurboID^fl/fl^”). F. Comparison of protein enrichment across tissues. G. Dynamic range plot for previously published neuronal and subtype markers in the DRG (LaCroix-Fralish et al., 2007; Tavares-Ferreira et al., 2022; Zheng et al., 2019), plotted as the median expression across all compartments.

### TurboID-mediated enrichment of proteins across compartments

In Adv^Cre^;TurboID^fl/fl^, proteins were considered enriched based on two criteria: A; proteins detected in Adv^Cre^;TurboID^fl/fl^ samples, but not in TurboID^fl/fl^ samples, or B; proteins significantly upregulated in Adv^Cre^;TurboID^fl/fl^ samples compared to TurboID^fl/fl^ (log_2_ fold change (LFC) > 1, FDR < 0.05,see Methods) (Fig 2D-F, STable 2). These upregulated proteins are referred to as differentially expressed proteins (DEPs). In addition, there are few downregulated proteins, likely reflecting endogenous biotinylation such as in the case of Acaca, Acacb, Mccc1, and Pcca (Fig 2D), which are routinely reported in proximity labelling studies (Dumrongprechachan et al., 2021; Sun et al., 2022).

All enriched proteins (based on criteria A and B above) were then combined, compared across compartments (Fig 2E-F), and treated equally for all downstream analyses. We identified the highest number of enriched proteins in the somata (DRG), followed by the central terminals (terminals_c_, SC), axons (SCN), and finally the peripheral terminals (terminals_p_, skin), with overlapping and distinct proteins (Fig 2E-F). Across the dynamic range, we see an enrichment of neuronal cell type markers (e.g. Calca, Osmr, Piezo2) and key nociceptor ion channels (e.g. Scn8a-10a, Trpv1) in Adv^Cre^;TurboID^fl/fl^ samples over TurboID^fl/fl^ (Fig 2G, SFig 2D; full list of proteins in STable2).

Of note, in the skin, Adv^Cre^ is known to be expressed in both primary afferents and, to a lesser extent, Merkel cells (Hunter et al., 2018). Because of this, we also detect Merkel cell markers (e.g. Krt20) in our TurboID-enriched proteome of the skin but not in other compartments (STable 2).

Taken together, our results yield a curated, enriched, and compartmental proteome of murine primary sensory neurons ranging from hundreds to thousands of proteins per compartment.

### Enrichment of proteins in neuronal terminals

A complementary method to isolate neuronal compartments from the skin or spinal cord is synaptosome isolation. Synaptosome preparations allow for the co-isolation of pre- and post-synaptic machinery, enriched against other components in the tissue.

Recently, synaptosomes were profiled from both mouse and human skin without tailoring for a specific cell type (Rawat et al., 2025). While the methods diverge in the exact tissue extracted, both our proximity labelling approach and synaptosome profiling from either species should extract the synaptic compartment of sensory terminals. To test this, we assessed overlapping proteins in both enriched datasets and compared cellular compartment pathways on this overlap (Fig 3A, mouse; Fig 3B, human). Indeed, we see a significant enrichment for synapse-related pathway terms across mouse datasets when comparing overlapping proteins (Fig 3A). There is also a trending enrichment when comparing our mouse TurboID data from peripheral terminals with human skin synaptosomes (Fig 3B, FDR ∼ 0.07-0.09). The latter likely reflects reported similarities in identified proteins despite known species differences (Segelcke et al., 2023). Overall, this comparison strengthens and validates our TurboID data in terminal endings and, at the same time, complements the published synaptosome data. While both TurboID and synaptosome profiling extracted synapse-related proteins (Fig. 3A), in our TurboID dataset we detect hundreds of proteins associated with synaptic compartments that were not reported in the paw synaptosome fraction (Rawat et al., 2025), such as Syn1, Syn2, and Synj1 (Fig. 3C, STable 3).

**Figure 3.**
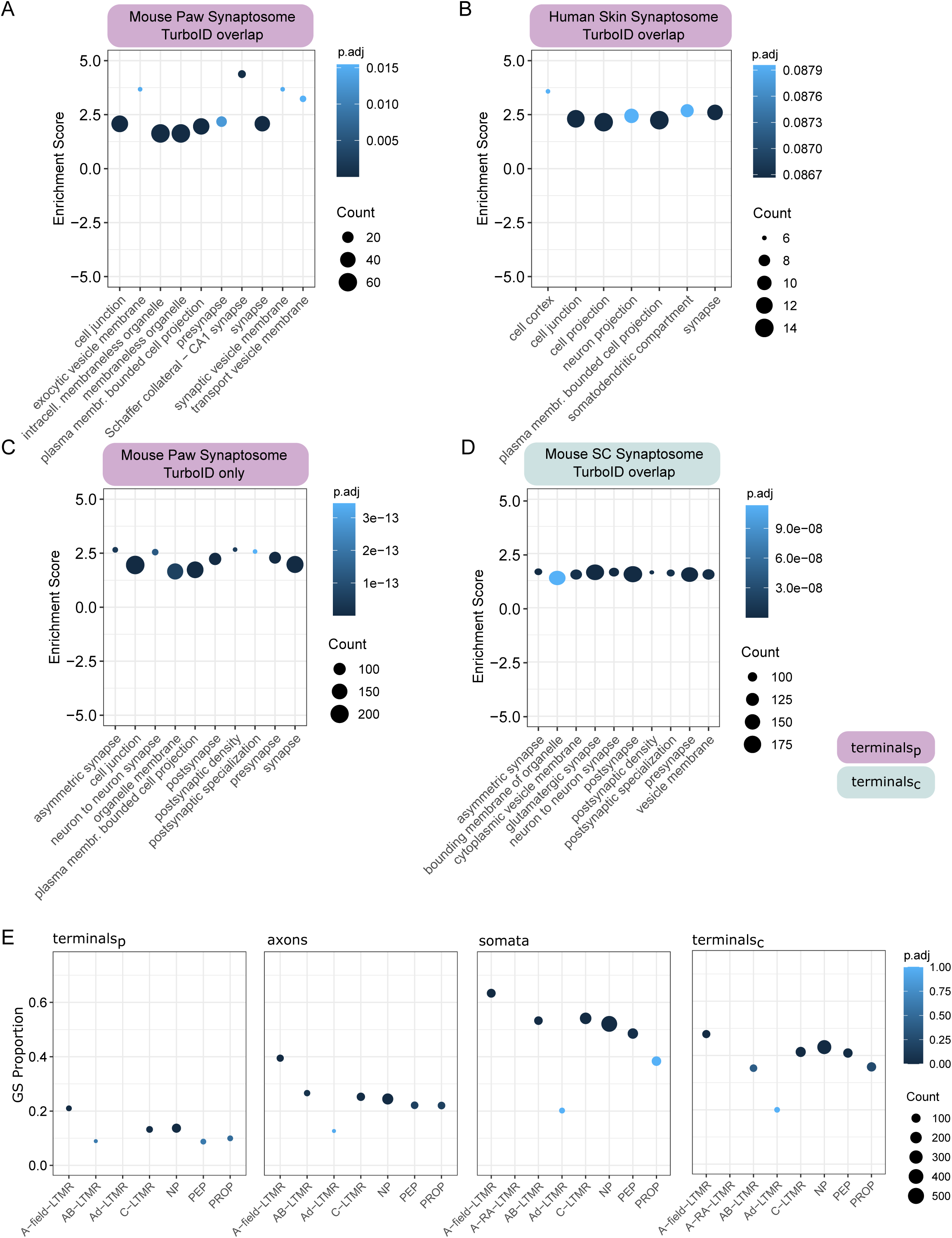
Validation against published datasets. A-C. Comparison of our TurboID proteome in peripheral terminals with Rawat et al., (2025) skin synaptosome proteomics, via overenrichment analysis for cellular compartment terms. A. Overlapping pathways with the murine synaptosome dataset. B. Overlapping pathways with the human skin synaptosome dataset. C. Proteins enriched in our TurboID peripheral terminal data but not detected in the murine synaptosome dataset. D. Comparison of our TurboID proteome in central terminals to spinal cord (SC) synaptosome data from Casañas & Montesinos (2022). E. Gene set enrichment analysis against custom DRG subtype gene lists. GS = gene set. NP = non-peptidergic. PEP = peptidergic. PROP = proprioceptor. LTMR = low threshold mechanoreceptor. LFC = log_2_ fold change. terminals_p_ = peripheral terminals (skin); terminals_c_ = central terminals (SC). Count = the number of proteins detected within each gene list, per compartment. Extended data in STable 3.

Next, we tested the overlap with a murine spinal cord synaptosome dataset (Casañas & Montesinos, 2022), which reports both first- and higher-order synapses from a diverse population of cell types (Fig 3D). Again, we see a positive enrichment of synapse terms across datasets further validating that our TurboID approach successfully extracts relevant synaptic compartments.

The use of Adv^Cre^ allows targeting the majority sensory neurons in the PNS (Zurborg et al., 2011). To evaluate the representation of neuronal subpopulations within our dataset, we performed gene set enrichment analysis using a curated list of neuronal subtype markers from published deep RNA-seq data (Barry et al., 2023). As expected, the strongest subtype enrichment was observed in the DRG, with significantly enriched signatures for non-peptidergic (NP) and peptidergic nociceptors (PEP), as well as a mix of low-threshold mechanoreceptors (LTMRs; including A-field-LTMR, C-LTMR, and AB-LTMR) (Fig 3E). Across all compartments a positive trend for neuronal subtypes emerged. Significant enrichments were particularly notable for NP nociceptors, A-field-LTMR, and C-LTMR (Fig 3E, STable 3). These findings indicate that our proximity labelling approach preserves the molecular diversity of sensory neuron subtypes across their anatomical domains.

### Compartmental sexual dimorphism

To explore sex-specific molecular differences, we analysed differential protein expression between male and female mice across all compartments (SFig 3). With an FDR < 0.05, the TurboID-enriched proteome appeared largely conserved between sexes, with a few exceptions in abs(LFC) > 1, such as Lgi1 and Sprr1a in central terminals and Brd4 and Palm2 in the peripheral terminals (SFig 3A, STable 4).

To gain broader biological insights, we also conducted gene set enrichment analysis using ranked LFC. This analysis revealed sexually dimorphic regulation of few pathways (SFig 3B). In the peripheral terminals, we see differences in circulatory and vascular pathways, whereas peptide biosynthesis and macromolecule biosynthesis show differences in the somata. We further explored potential sexual dimorphism through phenotype permutations as a more conservative approach and did not find any regulated pathways in any compartment. Taken together, our data support a narrative of minimal sexual dimorphism in TurboID-enriched proteomes.

### Functional signatures show separation by neuronal compartment

To extract the corresponding functional signature, Gene Ontology (GO) pathway analyses were performed against a background of all proteins expressed in the corresponding tissues (Fig 4). While neuronal signatures were detected in each compartment, they also exhibited functional specificity(Fig 4, STable 5). Primary afferents synapse on secondary order neurons in the dorsal horn of the spinal cord. This corresponds to the enrichment of proteins involved in presynaptic function and synaptic signalling between neurons, as well as vesicle-mediated transport, exocytosis, and other synapse-related terms in the SC. The DRG contain neuronal somata, and, thus, corresponding functional enrichment matches expectations. Here, we observed the enrichment of proteins involved in mRNA processing, post-translational modification, and nuclear structures, in line with the biosynthetic activity expected in neuronal cell bodies.

**Figure 4.**
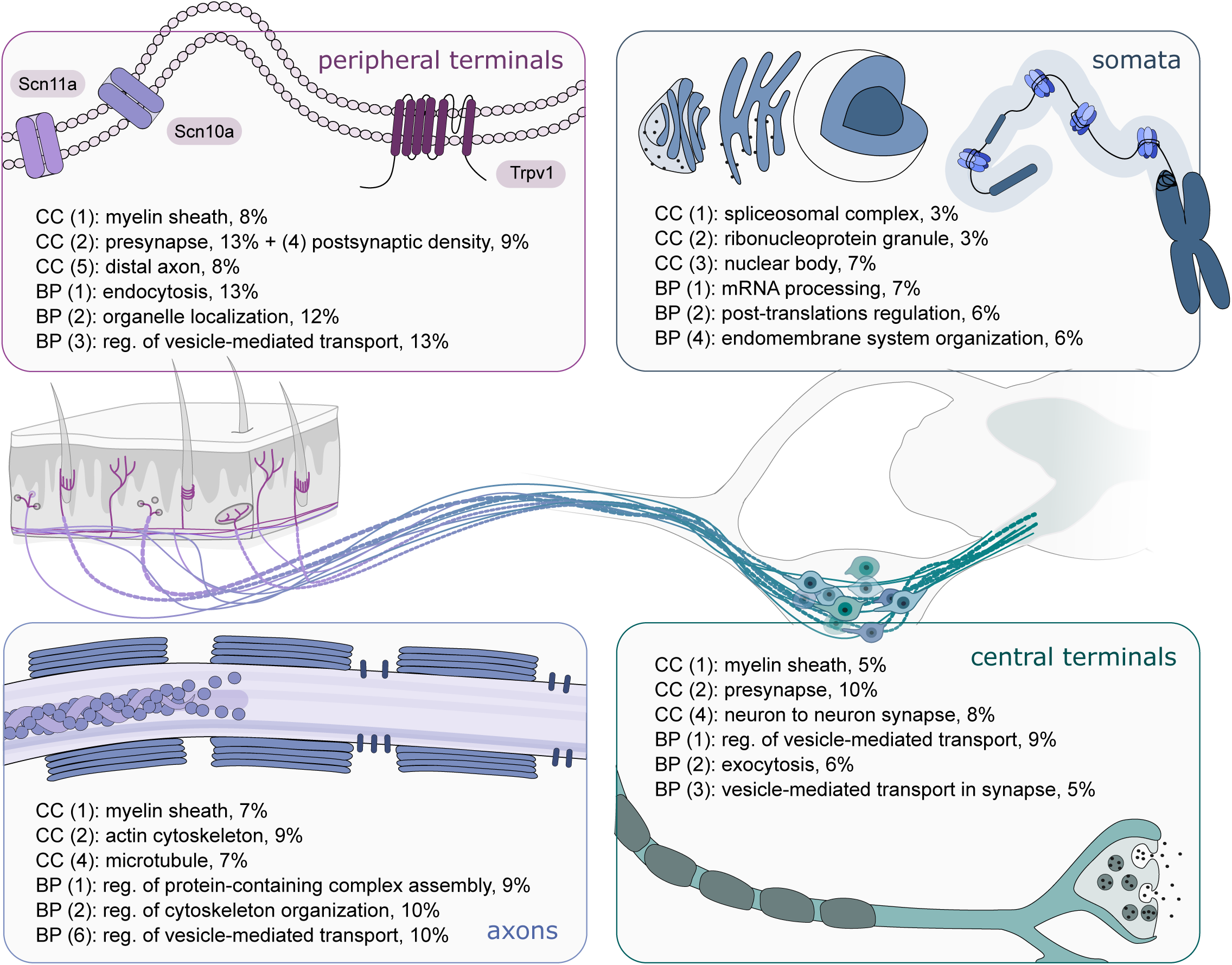
Pathway and neuronal subpopulation enrichment across compartments. Schematic highlights the GO over-enrichment analysis per tissue. BP = biological pathway; CC = cellular compartment. (X) term rank by adjusted p-value. % represents the GO term/neuronal enrichment. Please see STable 5 for corresponding data.

**Figure 5.**
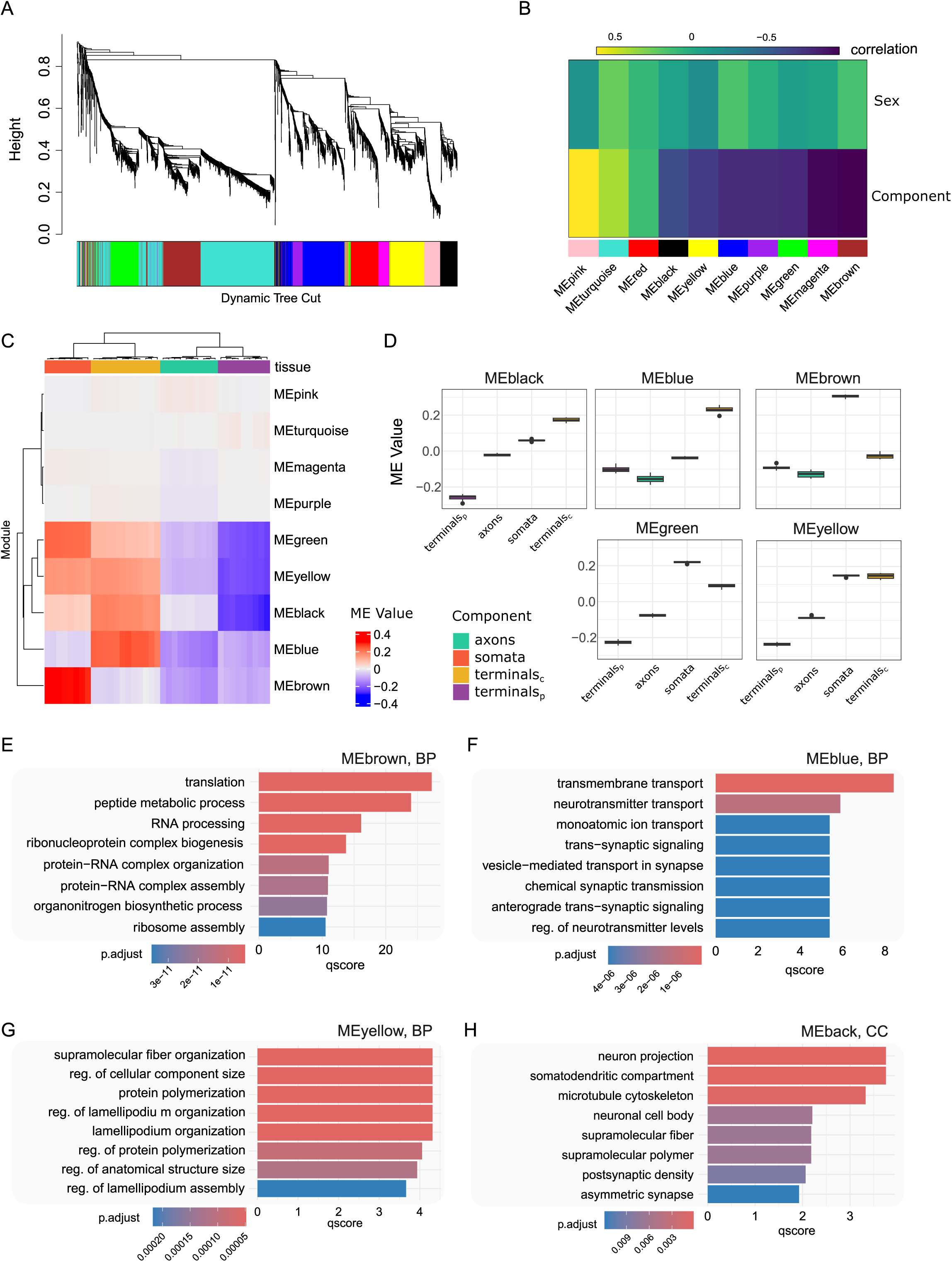
Weighted co-expression network of the TurboID-enriched proteome. A. Dendrogram with minimum 30 proteins per module. B. Correlation between modules and traits (compartment or sex). C. ME (module eigengene) value per sample. D. Module eigengene (ME) separation by component and module for modules significantly correlated to tissue (FDR < 0.05, BH correction). E-H. Biological pathway (E-G) and cellular compartment (H) analysis for modules of interest. E. brown, F. blue, G. yellow, H. black. No biological pathways or cellular components were significant for the green module. p.adjust = adjusted p-value. qscore = −log_10_(p.adjust). terminals_p_ = peripheral terminals (skin); terminals_c_ = central terminals (SC).

The sciatic nerve consists of neuronal axons, as well as Schwann cells, fibroblasts, endothelial cells, and immune cells. Its neuronal proteome is enriched for myelination, cytoskeletal organization, and vesicle-mediated transport (STable 5). The “myelination” terms included nodal and paranodal proteins such as Cntn1 (Contactin-1) and Cntnap1 (Caspr1), as well as more generally expressed neuronal proteins like Ncam1, Nefl, Nefm, and Nefh. In addition, we found proteins more classically associated with myelin itself, such as Plp1, MBP, and MPZ , which also show expression in neuronal subtypes (Bhuiyan et al., 2024).

To explore functional organization across neuronal compartments, a signed weighted protein co-expression network was constructed on the neuronal proteome (Fig 5). The co-expression network was built using the WGCNA package in R with a soft power threshold of 14. Ten protein co-expression modules were identified (excluding the unassigned “grey” module), each comprising a minimum of 30 proteins (Fig 5A). Of these, 9 modules showed significant correlation with anatomical compartment (FDR < 0.05), while none were associated with sex (Fig 5B). The latter is in line with our pathway analysis (SFig 3) and consistent with earlier findings reporting a mild sexual dimorphism in DRG neurons (Barry et al., 2023; Tavares-Ferreira et al., 2022).

Specific modules showed clear compartmental specificity. For example, the brown module correlated strongly with the DRG (somata), while the blue module was associated with central terminals in the spinal cord. Conversely, several modules -including black, yellow, and green - were negatively associated with peripheral terminals, likely indicating a relative depletion of these proteins in distal axonal endings (Fig 5C–D). As expected, most modules exhibited strong alignment with one or more anatomical compartments, reinforcing the functional specialization of the sensory neuron proteome across space (Fig 5D).

Biological pathway analyses of these modules (against a background of all proteins used to build the co-expression network) was then performed (Fig 5E-H, STable 6). The ‘brown’ module, which is positively correlated with the soma, reflects a group of proteins implicated in ‘translation‘, ‘peptide metabolic process‘, and ‘RNA processing‘(Fig 5E). The blue module, which correlated with central terminals in the spinal cord, showed associated with ‘trans-synaptic signalling‘, ‘regulation of neurotransmitter levels‘, and ‘neurotransmitter transport‘, reflecting synaptic functions in the dorsal horn (Fig 5F). The yellow module, which was positively correlated with soma and central terminals, relates to ‘protein polymerization‘, ‘supramolecular fibre organization’ and the ‘regulation of cellular component sizè(Fig 5G).

No modules were positively correlated with the peripheral terminals - possibly due to the lower number of enriched proteins in peripheral terminals - but the most negative correlation is the black module. Although no biological pathways reached statistical significance in this module, cellular components related to ‘neuronal projection‘, ‘somatodendritic compartment‘, ‘postsynaptic density‘, and ‘neuronal cell body’ were among the annotations (Fig 5H).

### TurboID proteomics provides a unique look at terminal endings

To explore protein interaction and functional organization in peripheral and central terminals, large-scale protein–protein interaction (PPI) networks were constructed using STRING (Fig 6, SFig 4) (Szklarczyk et al., 2023). A Girvan-Newman fast greedy algorithm was then used to generate community clusters (Su et al., 2010; Utriainen & Morris, 2023), and the functional enrichment of each cluster was calculated.

**Figure 6.**
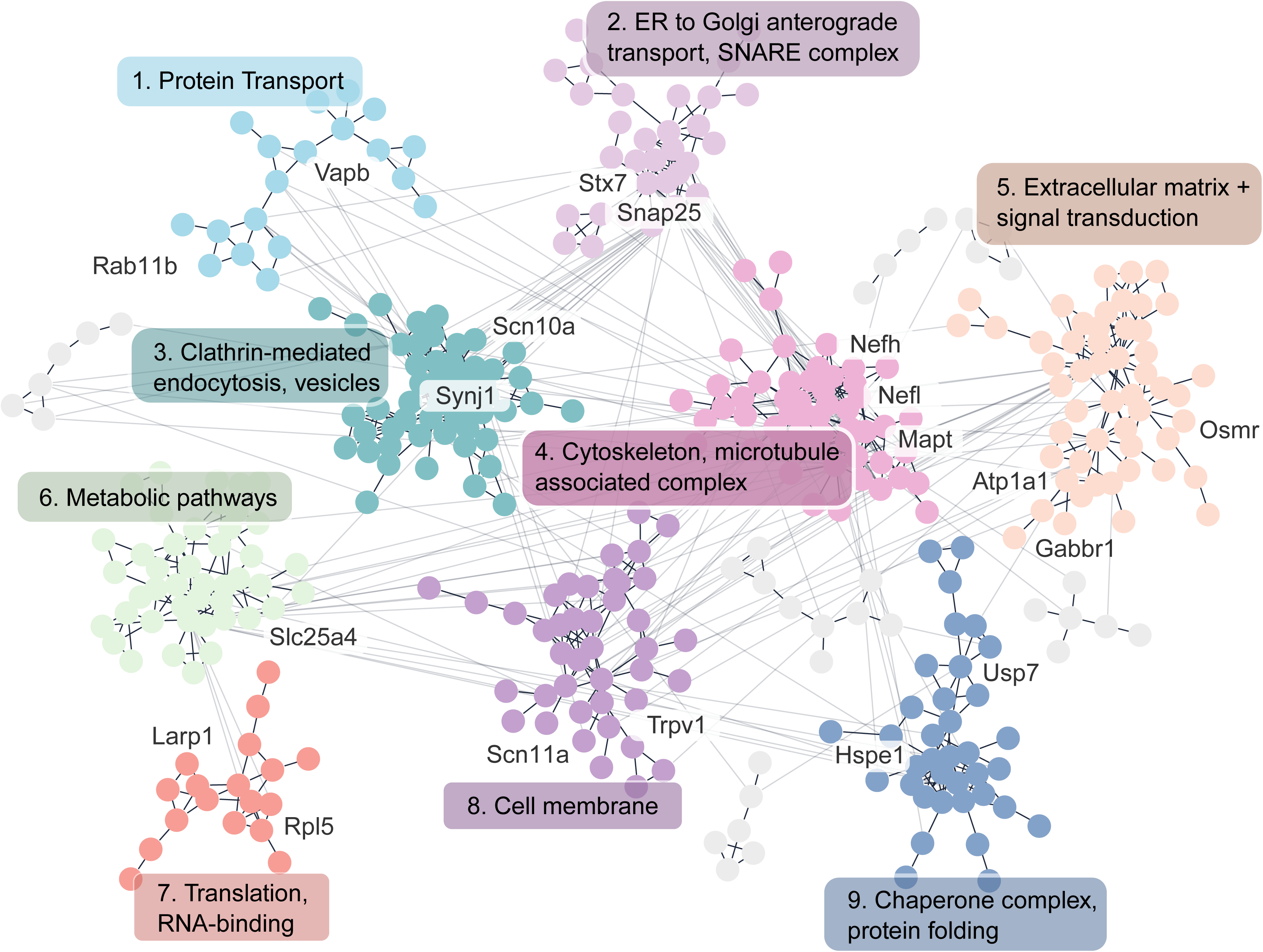
Community clustering of the peripheral terminal proteome, with edges denoting high confidence protein-protein interactions. Protein nodes are coloured by cluster, and clusters are described by functional summaries with proteins of interest highlighted. Full functional enrichments are available in STable 7.

In the peripheral terminals (skin), the proteome revealed a diverse and functionally rich landscape, spanning protein transport, metabolic pathways, and signal transduction (Fig 6). A wide array of protein classes was detected, including SNARE complex components such as Syntaxin-7 and Snap25, as well as chaperones and ubiquitin-related proteins, such as Usp7 and heat shock proteins. Notably, even pain-associated ion channels (Trpv1, Scn10a, and Scn11a) were detected in clusters annotated for membrane trafficking and exocytosis. A full breakdown of each cluster enrichment is available in STable 7.

A similar pattern was observed in the central terminals of the spinal cord (SC), where clustering revealed extensive functional heterogeneity (SFig 4, STable 8). Enriched clusters included pathways involved in signal transduction, endosomal processing, and membrane trafficking—hallmark processes of synaptic signalling and neurotransmission (see Stable 8 for a functional breakdown).

Together, these analyses demonstrate that proximity labelling via TurboID enables unprecedented insights to the proteomic architecture of otherwise inaccessible neuronal compartments. This approach holds considerable promise for extracting functional, spatially resolved proteomes in defined cell types.

### TurboID proteomics enables deeper insights than bulk tissue proteomes

To contextualize the advantages of TurboID-based proximity labelling, we compared our compartment-resolved proteomes to conventional bulk tissue proteomics. We harvested the skin, SCN, DRG and SC and determined the proteome using the same mass spectrometry-based workflow as applied to the TurboID samples above (SFig 5, STable 9). As expected (Barry et al., 2018; Xian et al., 2022), we detect several thousands (7000-9000) protein groups per tissue (SFig 5A), with samples clustering by tissue (SFig 5B).

Consistent with earlier studies (Barry et al., 2018; Xian et al., 2022), we identified many neuronal proteins in bulk datasets. While these bulk tissue proteomes exhibited a large overlap with TurboID-enriched proteomes, the latter identified dozens of additional proteins in each compartment (STable 10; example for bulk skin versus TurboID-enriched peripheral terminals, SFig 5D).

To additionally assess the enrichment achieved by TurboID proteomes, we compared the extent of immune cell subtypes present in TurboID proteomes versus bulk tissue proteomes. (SFig 5E). ImmuCellAI (Miao et al., 2022) was used to calculate an “immune infiltration score”, which reflects a composite GSEA enrichment across immune cell subtypes. Indeed, compared to bulk tissue samples, we obtained a reduced immune signature in TurboID-enriched proteomes, further validating our approach (SFig 5E).

This highlights two key advantages of TurboID-based proteomics: First, it facilitates the enrichment of proteins in a Cre-dependent manner within the same tissue. Second, TurboID enhanced detection sensitivity in tissues with sparse cellular content, such as peripheral terminals in the skin. In these regions, the detection of sensory neuron-specific proteins — particularly ion channels — has been challenging and largely unsuccessful using bulk proteomics (Dyring-Andersen et al., 2020; Xian et al., 2022). In contrast, TurboID-based profiling recovered functionally important signatures in peripheral terminals, including prominent drug targets such as Scn10a, Scn11a, and Trpv1, providing molecular access to previously inaccessible compartments (Fig 6, SFig 5D).

Together, these data underscore the power of TurboID-based proteomics to resolve cell type-specific proteomes within complex tissues. Importantly, this strategy opens a path to study the proteome dynamics of afferent terminals in disease models and in response to pharmacological interventions.

### TurboID proteomics reveals neuron-enriched changes in response to oxaliplatin treatment

Next, we tested the utility of our TurboID workflow in a pharmacological context. For this, we turned to oxaliplatin, a chemotherapeutic agent. Chemotherapy-induced peripheral neuropathy (CIPN) is a common and often painful condition affecting the peripheral nervous system, known to impact neuronal and non-neuronal cells in the DRG (Sisignano et al., 2014). In adherence to the 3Rs of animal research, we employed a DRG explant model to explore oxaliplatin-induced changes in neuronal somata (Fig 7, SFig 6-7, STables 11-16). Mirroring the compartmental proteomes above, we determined the bulk proteome of DRG explants from wildtype mice and, thereafter, compared these data with TurboID-enriched proteomes obtained from Adv^Cre^;TurboID^fl/fl^ mice (versus TurboID^fl/fl^ controls).

**Figure 7.**
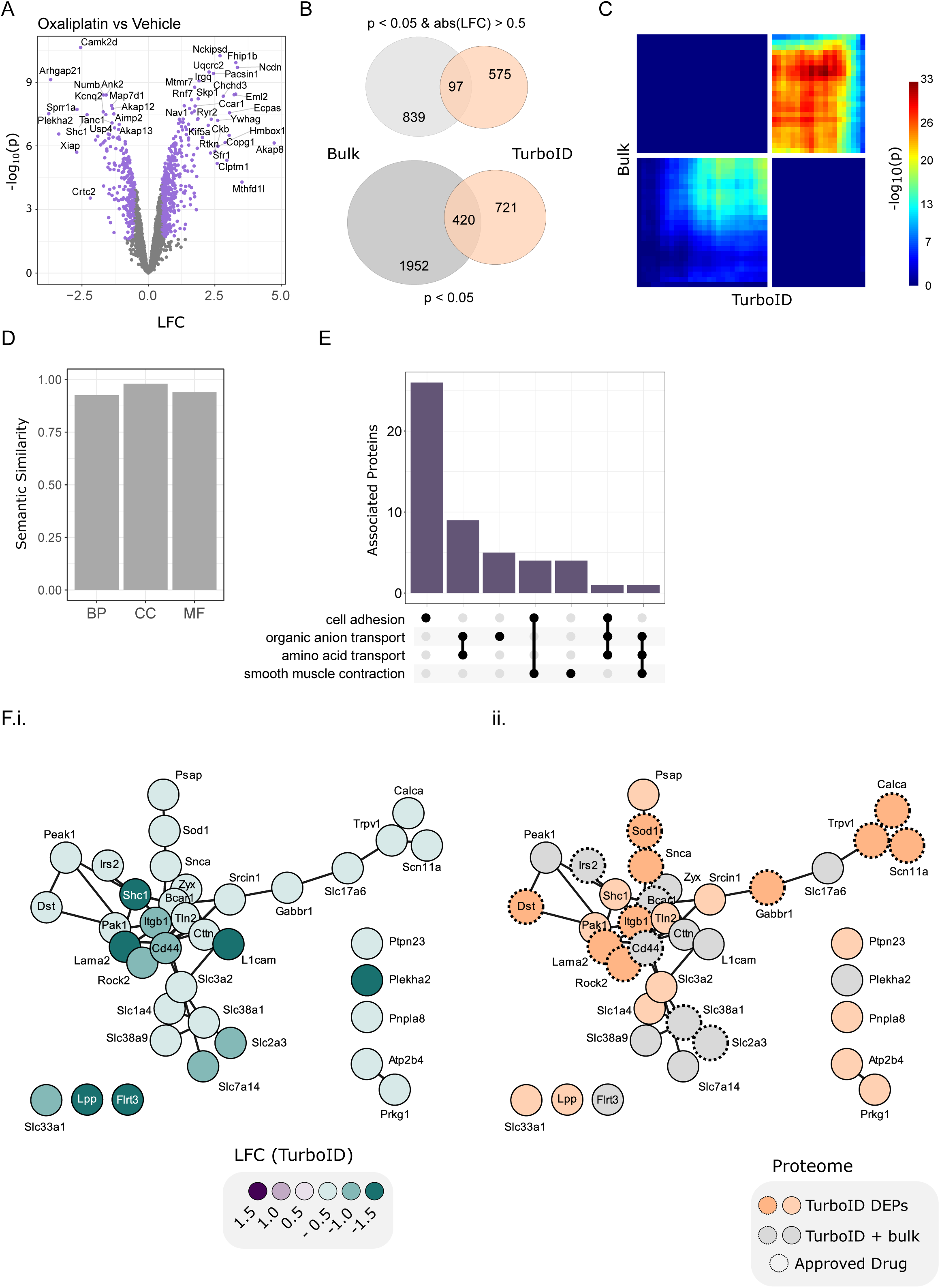
Oxaliplatin-treated Adv^Cre^;TurboID^fl/fl^ explants highlight neuronal dysregulation. A. Volcano plot of the TurboID proteome (Adv^Cre^;TurboID^fl/fl^) for oxaliplatin vs vehicle,. B. DEP overlap (oxaliplatin vs vehicle) between bulk (WT, grey) and TurboID (orange) explant proteomes. C. Correlation between LFC for overlapping DEPs in B. D. Semantic similarity between TurboID and bulk DEPs. E. Neuronal biological pathway downregulation after oxaliplatin treatment, highlighting protein counts which overlap across pathways. F. STRING protein-protein interaction network for DEPs in regulated pathways from E. i, LFC in the TurboID-enriched proteome. ii, Approved drug targets from dgidb.org are highlighted with dashed circles. DEPs detected only in the TurboID-enriched proteome are in orange, those overlapping with the bulk analysis are in grey. LFC, log2-fold change.

In the bulk proteome of wildtype mice we identified ∼7200 protein groups (SFig 6A, STable 11), with clustering driven by oxaliplatin concentration (SFig 6B-C). Consistent with earlier studies, oxaliplatin exposure led to widespread proteome changes, including alterations in known neuronal proteins (SFig 6D-E, STable 12). When comparing DEPs from our DRG explant data to previous work *in vivo* (Yang et al., 2022), we see a strong correlation of protein expression (SFig 6F). We next employed a rank-rank hypergeometric overlap test and observed good concordance for DEPs across both datasets (SFig 6G). These comparisons suggest similar oxaliplatin-induced proteome changes validating our explant model.

Next, we treated DRG explants from Adv^Cre^;TurboID^fl/fl^ mice and TurboID^fl/fl^ controls with oxaliplatin or its vehicle (Fig 7, SFig 7, STable 13). As expected, Adv^Cre^;TurboID^fl/fl^ explants exhibited prominent protein enrichment compared to TurboID^fl/fl^ controls (SFig 7A). Samples clustered by genetics in the first principal component (PC), with slight separation by drug treatment in the second PC (SFig 7B). In both oxaliplatin-treated (SFig 7C, left) and vehicle-treated (SFig 7C, right) explants, Adv^Cre^;TurboID^fl/fl^ samples (vs TurboID^fl/fl^) showed robust TurboID-mediated enrichment with ∼ 2500 protein groups per condition (SFig 7D, STable 14).

Within this TurboID-enriched proteome, we detected a substantial number of DEPs after oxaliplatin treatment compared to vehicle-treated controls (Fig 7A, STable 15). Notably, these differences were only partially reflected in the bulk explant proteome (Fig 7B).

Even so, we saw a high rank-rank hypergeometric overlap between the bulk explant and TurboID-enriched proteomes after oxaliplatin treatment (Fig 7C), and a high semantic similarity across DEPs (Fig 7D).

Next, we assessed DEP-associated functions in DRG explant proteomes. After oxaliplatin treatment, pathways related to amino acid transport and cell adhesion were downregulated in the TurboID-enriched proteome (Fig 7E, STable 16). To further dissect these pathways, we constructed a corresponding PPI network based on pathway-associated DEPs (Fig 7F). This revealed prominent downregulation across multiple functional modules (Fig 7Fi). While some of these DEPs were regulated in both bulk and Tur-boID-enriched explant proteomes, others were only detected in the TurboID-enriched proteome (Fig 7Fii). The latter included Trpv1, Scn11a, and Calca, as well as the central module with Itgb1, Lama2, and Rock2 – all of which are targets of approved drugs (https://dgidb.org/). For example, capsaicin patches (target: Trpv1) for pain, and the kinase inhibitor Vandetanib (target: Rock2), which is prescribed against a subset of thyroid cancers (Viola et al., 2016) (https://dgidb.org/).

These findings illustrate the enhanced resolution of TurboID-mediated proteomics, uncovering molecular changes that may be masked in bulk tissue analyses. Thus, our TurboID workflow offers a powerful tool to dissect disease mechanisms and treatment effects in complex tissues.

## Discussion

As a generalized proof-of-concept, we demonstrate how our TurboID (TurboID^fl/fl^) mouse line enables cell-type-specific proteomic labelling across anatomically distinct compartments. We paired it with Adv^Cre^ to target TurboID expression to primary sensory neurons of DRG, resulting in robust biotinylation across neuronal compartments. Using DIA-PASEF mass spectrometry, we provide unprecedented insights into compartment-specific DRG neuron proteomes from peripheral to central terminals.

This spatial resolution is particularly significant in the peripheral terminals of DRG neurons in the skin. Peripheral nerve endings are typically characterized by immunohistochemistry and have remained largely inaccessible to unbiased omics approaches. Neither our bulk proteomic data nor previous high-coverage DIA-MS datasets from mouse skin or human skin were able to profile key sensory neuron markers (Dyring-Andersen et al., 2020; Rawat et al., 2025; Xian et al., 2022). Sparse molecular signatures are often masked by dominant cell types, likely restricting the detection of neuronal signatures in the skin. Similar considerations apply to transcriptome profiling. While single-cell and single-nucleus RNA sequencing are valuable for cell-type-resolved transcriptomes, they are typically limited to somata within the DRG, lack proteomic resolution, and do not capture distal compartments (Grindberg et al., 2013; Matzinger et al., 2023). The expansion of spatial sequencing helps address many of these issues, although recent spatial transcriptomics of the sciatic nerve have not resolved neuron-specific clusters (Ferreira et al., 2024), and to our knowledge, a high resolution spatial-seq dataset including peripheral terminals in the skin is still missing. Here, we used TurboID-based proteomics and were able to profile neuronal markers and prominent drug targets such as Trpv1, Scn10a, and Scn11a in the peripheral terminals. Consequently, at present, our TurboID-based proteomic atlas offers a unique spatial profile spanning the extent of murine DRG sensory neuron compartments. We expect that our data will provide a valuable protein-level stepping stone for ongoing efforts to expand spatial -omics to previously inaccessible cell types and compartments.

A strength of the TurboID approach is the ability to probe proteins enriched in one compartment over others. Many of the proteins enriched in central terminals relate to synapse function and vesicle trafficking as expected. Alongside, we also detected several proteins relevant to pain and somatosensation such as Lgi1, Stoml3, and Cacng2 (stargazin) (Bortsov et al., 2019; Farah et al., 2024; Lafrenière & Rouleau, 2011; Wetzel et al., 2017). In the axon, we a number of functionally relevant proteins were enriched, including Arhgap32, Atp1a2, and Kcnab3. Arhgap32 (aka Grit, Rics, or p250GAP) is a GTPase regulator that interacts with Ntrk1 to regulate neurite outgrowth (Kannan et al., 2012; Nakamura et al., 2002). Atp1a2 encodes an ATPase Na+/K+ transporting subunit with a role in migraine (Murata et al., 2020), and Kcnab3 is a shaker-type potassium axillary unit, likely expressed in the paranodal and juxtaparanodal region (Rasband & Trimmer, 2001). Shaker-type channels are general involved in regulating excitability (Rasband et al., 2001), but the implications of Kcnab3 expression remains unclear and follow-up work for all candidates is warranted for a better understanding of DRG sensory neuron functions.

While not a main focus of this study, we were also able to probe neuronal sex differences across compartments, expanding on previously reported sexual dimorphism in mouse and humans (Barry et al., 2023; North et al., 2019; Tavares-Ferreira et al., 2022; Xian et al., 2022). Mirroring these data across sensory neuron subtypes, we see marginal differences between male and female neurons under naïve conditions.

To highlight how TurboID proximity labelling could be exploited to study functional changes in a clinically-relevant disease context, we investigated proteome dynamics in a DRG explant model of CIPN. Exposure to oxaliplatin resulted in widespread proteomic alterations in both bulk and TurboID-enriched proteomes, as expected. While DEP sets varied between bulk and TurboID enrichments, semantic similarity of enriched pathways was high, indicating shared biological responses. A major benefit of our TurboID approach in explants is the neuronal enrichment, highlighting disruption to organic anion and amino acid transport pathways. This contrasts the more generalized effect of oxaliplatin seen in the whole DRG proteome composed of diverse cell types and highlights the practicality of the DRG explant model for rapid *in vitro* screening of neurotoxic effects.

Transporter disruptions have widely been reported in the context of CIPN (Filipski et al., 2009; Huang et al., 2020; Litke et al., 2022; Schulte & Ho, 2019; Sisignano et al., 2014; Sprowl et al., 2013). Organic cation transporters, such as Oct2, have been implicated in oxaliplatin uptake by sensory neurons, and their activity contributes to neurotoxicity (Sisignano et al., 2014; Sprowl et al., 2013). Similarly, organic anion-transporting polypeptides (OATPs) affect CIPN pathophysiology (Leskelä et al., 2011; Li et al., 2023; Schulte & Ho, 2019). Impaired transporter function can lead to drug accumulation in DRG, contributing to mitochondrial dysfunction, oxidative stress, and neuroinflammation, all of which underlie CIPN pathogenesis (Sisignano et al., 2014). In our study, several transporters were downregulated in response to oxaliplatin treatment. Some changes are evident across neuronal and bulk proteomes (e.g. Slc7a14, Slc2a3), while the dysregulation of others (e.g. Slc1a4, Slc3a2, Slc33a1) could only be detected when studying the neuronal proteome. Pursuing these transport pathways may provide novel strategies to mitigate CIPN while preserving the anticancer efficacy of chemotherapy, with promising attempts to target transporters already in development (Li et al., 2023).

When paired with conventional bulk analyses, our approach offers a multi-layered view of the proteomic landscape in response to drug treatment and in disease models, complementing and extending *in vivo* work.

### Limitations

This study provides a quantitative, compartmental proteomic atlas of DRG sensory neurons and serves as a valuable reference for the field. However, it is not exhaustive, and reports a neuronal-enriched proteome (opposed to neuronal-specific). Particularly, the detection of low-abundance transmembrane proteins is known to be challenging. It may be further improved through advances in mass spectrometry sensitivity, biotinylation efficiency, and sample preparation protocols. Fewer protein identifications were obtained in peripheral terminals and axons - a likely reflection of heterogenous tissue composition and sparse neuronal content - resulting in a relative bias toward somata and central terminals, where neuronal protein density is higher. While our results represent a robust starting point, future studies with deeper coverage and within-compartment comparisons under various biological conditions will be critical for advancing our knowledge about the molecular features of terminal regions. Likewise, deeper coverage and higher sensitivity will enable future work on differentiating sensory neuron subtypes and their sparse peripheral terminals.

### Future directions

A key advantage of using TurboID transgenic mice is the opportunity to interrogate otherwise inaccessible cellular compartments. Prior work in both mice and humans has shown alterations in intraepidermal nerve fibre density across painful conditions (Middleton et al., 2021; Schmidt et al., 2022). Pairing our compartmental proteome with *in vivo* functional testing will provide a rich understanding of the underlying molecular changes and their correlation with behavioural outcomes in future studies.

Another major benefit lies in the flexibility of Cre-based recombination. Pseudobulk analyses are frequently used in scRNA-seq (and spatial) transcriptomics to compare within cell-type changes across conditions. Paired to adequate transgenic drivers and downstream DIA-MS, TurboID^fl/fl^ can be used to generate a corresponding proteome for multi-omic analyses within cell types. In the context of sensory neurons, this can be subtype targeting (e.g. Mrgprd^Cre^ or Th^Cre^) or injury-specific targeting (e.g. Atf3^Cre^) (Cooper et al., 2024; Zheng et al., 2019). More broadly, our approach could be extended to diverse cell types, such as fibroblasts in the skin, satellite glia in the DRG, or interneurons in the dorsal horn (to name a few).

## Conclusions

In this study, we generated a TurboID^fl/fl^ transgenic mouse, using it to systematically profile the proteomes of DRG sensory neuron compartments. Spanning from peripheral to central terminals, we provide a proteomic atlas across DRG sensory neuron compartments. Our findings reveal compartment-specific protein distributions that reflect specialized neuronal functions. Furthermore, we report previously unknown changes in an explant model of chemotherapy-induced peripheral neuropathy. Taken together, our methodological workflow and obtained data serve as a highly valuable resource for studying neuronal specialization and proteome dynamics in the peripheral nervous system.

## Code Availability

All code is available at https://github.com/aliibarry/turboID.

## Data Availability

The mass spectrometry proteomics data, including metadata, fasta, and gene group matrices for each dataset have been deposited to the ProteomeXchange Consortium via the PRIDE(Perez-Riverol et al., 2022) partner repository with the dataset identifier PXD062198. Processed data are also available on github (https://github.com/aliib-arry/turboID/data/).

For queries regarding the TurboID mouse line please contact Nils Brose or Noa Lipstein.

## Acknowledgements

The authors would like to thank Christiane Harenberg, Dayana Warnecke and Mia Wzietek from the AGCTLab (Department of Molecular Neurobiology, Max Planck Institute, MPI, for Multidisciplinary Sciences, Goettingen) for expert technical assistance and genotyping, as well as Nicole Kanta and Sabrina Grundtner (University of Vienna) for their excellent assistance. We are grateful to the staff of the animal house of the MPI and of the mouse house at the Division of Pharmacy (University of Vienna) for expert mouse husbandry and support. We further thank Dr. Kimmina, Dr. Schraepler, Dr. Fuenfschilling and Dr. Papadopoulos (all at MPI) for their support in all mouse-related issues, as well as Elham Barkhordar and Dr. Dominic Winter (University of Bonn, Germany) for initial discussions about the BioID approach. We thank all members of the Schmidt laboratory for fruitful discussions.

This work was funded in part by the Austrian Science Fund (FWF) [10.55776/P36554] (to MS), the University of Vienna (to MS), the Max Planck Society, the German Research Foundation, Excellence Strategy EXC-2049−390688087 (to NL), and CRC 1286 ″Quantitative Synaptology″, project A11 (to NL) and A09 (to NB).

## Author Contributions

Method optimization, sample preparation and data collection were performed by JRS with input from FX, DVG, and MS. Analyses were performed by AMB, with input from JRS, FX, and MS. Mass spectrometry was performed by FX with input from JRS and DGV. Conceptualization and funding acquisition by MS with input from DGV. Oxaliplatin experiments were performed by JRS and TH. TurboID^fl/fl^ mice were conceived, designed, generated and validated by NB, FB, DS, and NL. Manuscript was written by AMB, JRS, and MS with input from all authors. All authors approved the submitted version.

## Supplemental Figures

**Supplemental Figure 1.**
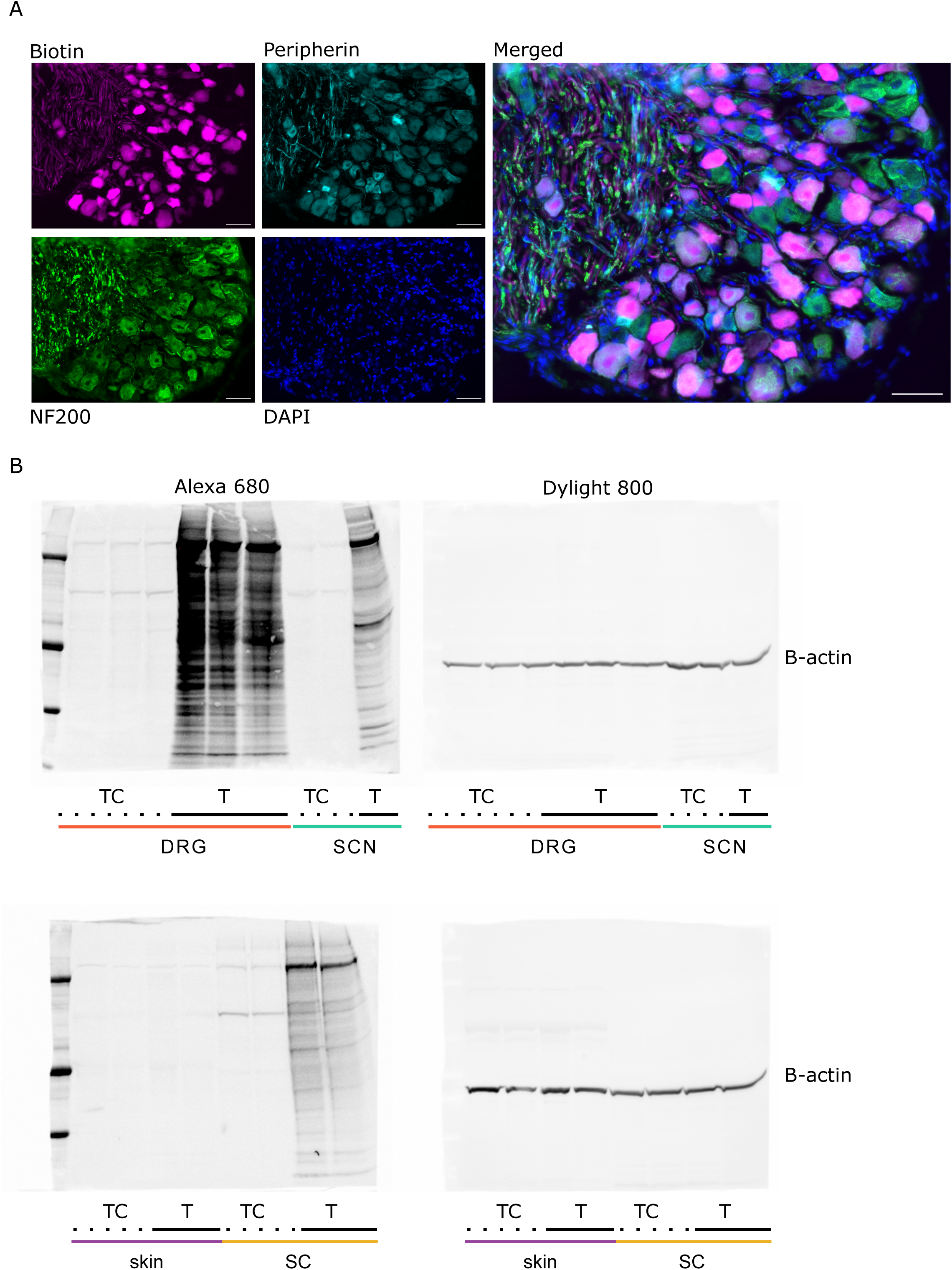
A. Immunohistochemistry of Adv^Cre^;TurboID^fl/fl^ with sensory neuron markers peripherin and NF200. Scale bar = 50 um. B. Western Blot analyses for TurboID (left) and B-actin loading controls (right). Adv^Cre^;TurboID^fl/fl^ (T) and TurboID^fl/fl^ (TC) samples for each tissue were loaded. DRG, dorsal root ganglia; SCN, sciatic nerve; SC, lumbar spinal cord.

**Supplemental Figure 2.**
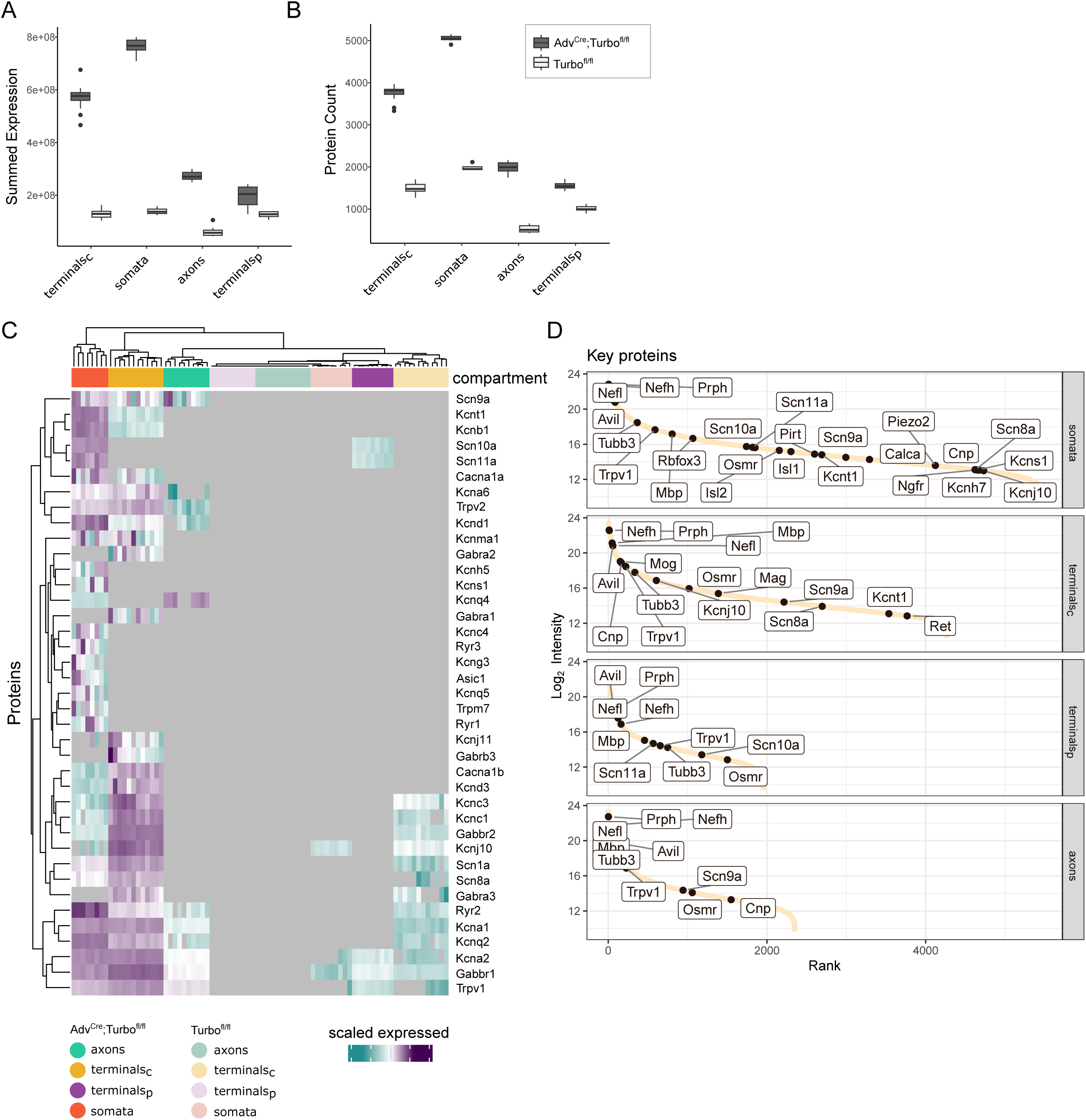
DIA-PASEF mass spectrometry of Adv^Cre^;TurboID^fl/fl^ and TurboID^fl/fl^ control samples across sensory neuron compartments. A-B. Summed expression (A), and Protein Group counts (B) by tissue and TurboID status. C. Heatmap of ion channel expression across all samples. D. Dynamic range for proteins of interest in Adv^Cre^;TurboID^fl/fl^ samples plotted as expression (log_2_ Intensity) per component. terminals_p_ = peripheral terminals (skin); terminals_c_ = central terminals (SC). Relates to Figure 2.

**Supplemental Figure 3.**
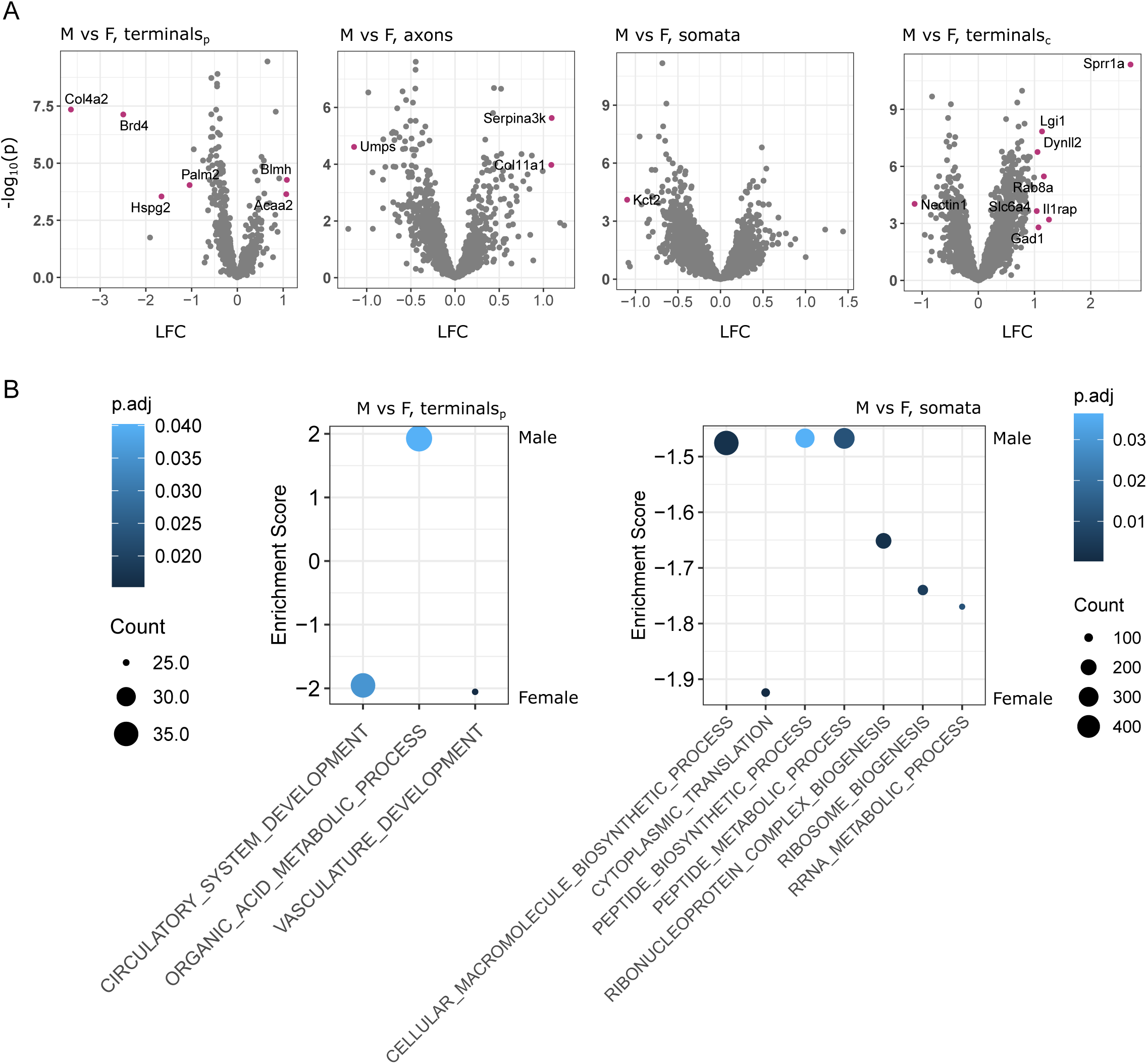
Sexual dimorphism across neuronal compartments. A. Differential protein expression by compartment, comparing male vs female mice within TurboID-enriched proteomes. Significant proteins are in pink. LFC = log_2_ fold change B. Biological Pathway analysis for peripheral terminals and DRG on ranked fold changes between male and female samples. Counts reflect the number of proteins in each respective term. terminals_p_ = peripheral terminals; terminals_c_ = central terminals.

**Supplemental Figure 4.**
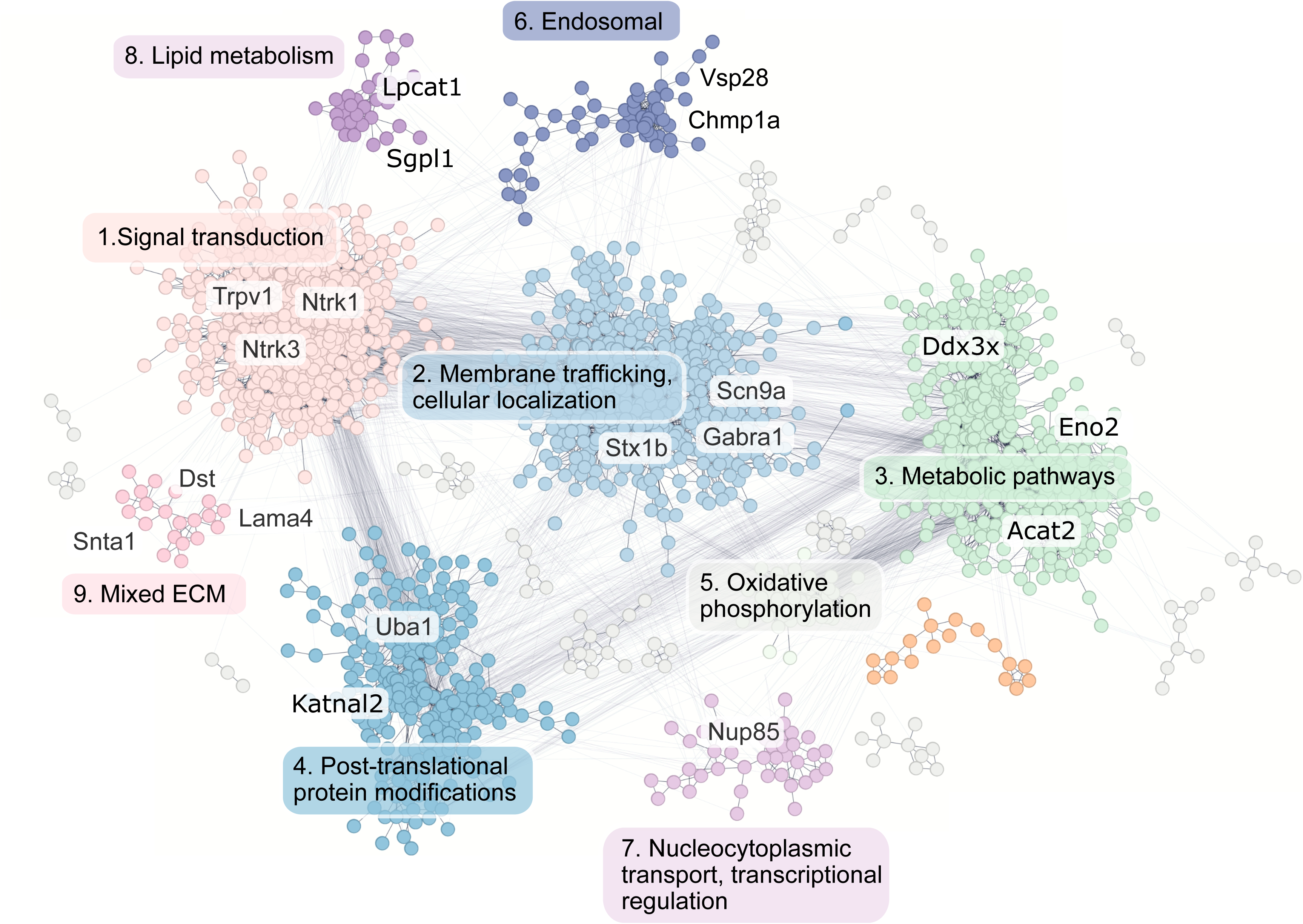
Community clustering of the TurboID-enriched central terminal proteome, with edges denoting high confidence protein-protein interactions. Protein nodes are coloured by cluster, and clusters are described by functional summaries with proteins of interest highlighted. Full functional enrichments are available in Stable 8.

**Supplemental Figure 5.**
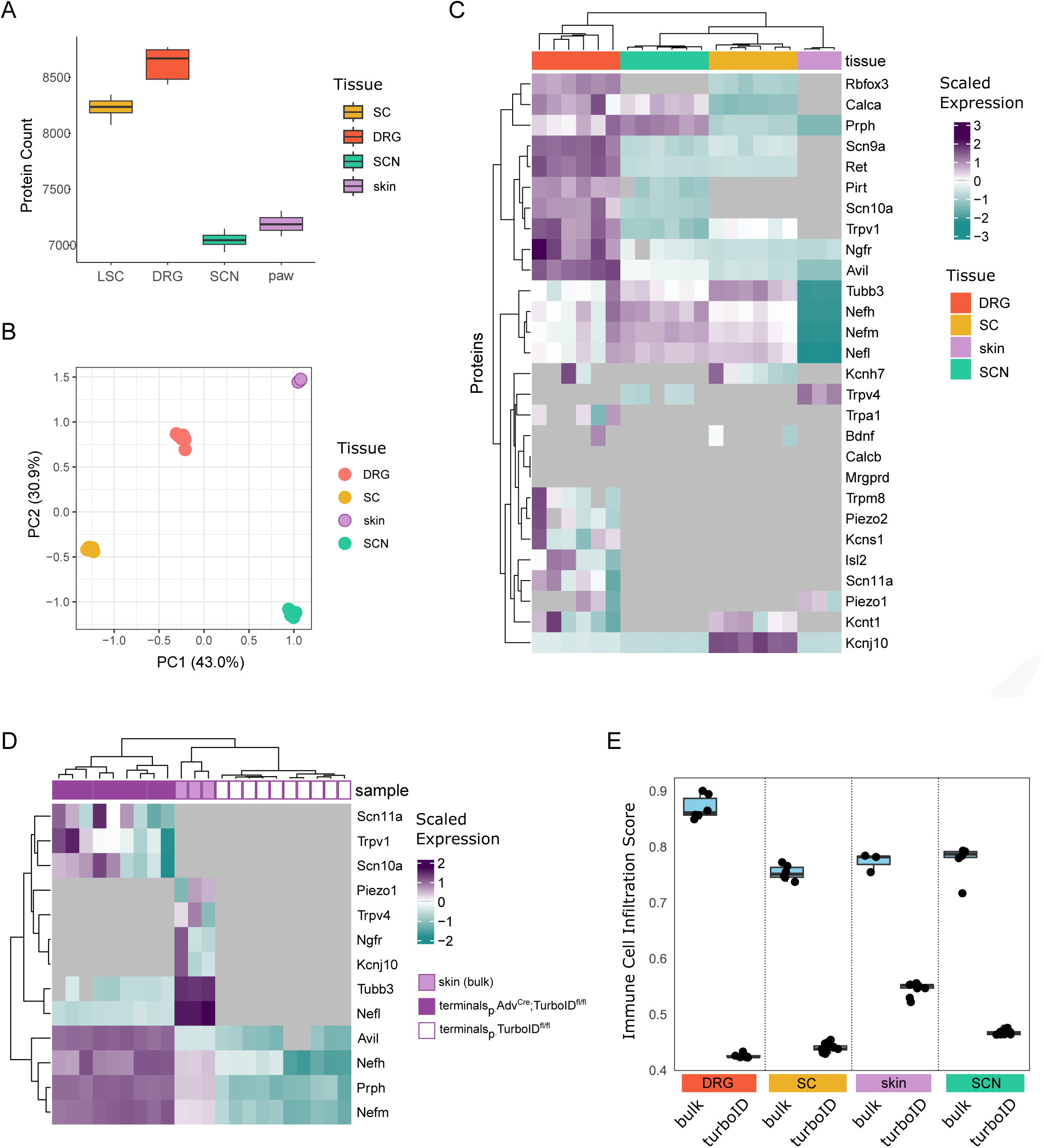
Bulk proteomes of the four analysed tissues from wildtype mice. A. Protein Group (PG) counts per tissue. B. PCA by tissue. C. Heatmap of the scaled expression of selected key neuronal genes. D. Comparison of key neuronal proteins between peripheral terminals of Adv^Cre^;TurboID^fl/fl^, those of TurboID^fl/fl^, and bulk proteomes obtained from skin samples. E. Depletion of ImmuCellAI immune cell signatures is evident in Adv^Cre^;Turbo^fl/fl^ samples compared to corresponding bulk samples from wildtype tissues.

**Supplemental Figure 6.**
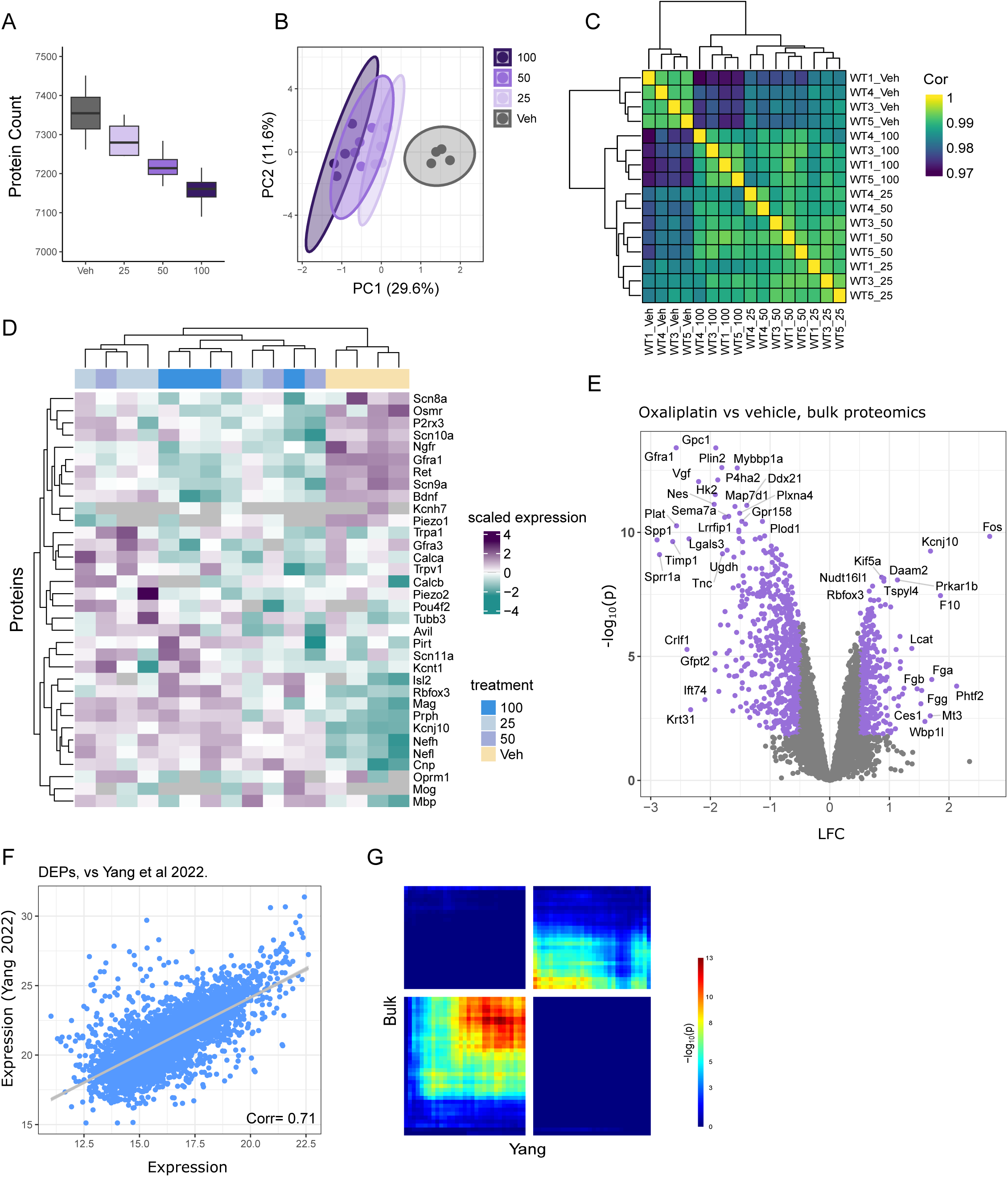
Bulk Proteomics of DRG explants exposed to oxaliplatin (25, 50 or 100 uM) or vehicle (Veh). A. Protein counts. B. PCA across sample conditions shows clear separation of oxaliplatin-treated samples versus Veh-treatment. C. Correlation across samples, labelled as oxaliplatin (25, 50, 100) or vehicle (Veh). D. Heatmap across selected key neuronal proteins. E. Differentially expressed proteins (DEPs) between 100 uM oxaliplatin- and Vehicle-treated DRG explants using bulk proteomics. Purple dots indicate significantly regulated proteins (FDR < 0.05, LFC > 0.5). F. Correlation of relative protein abundance between our DEP dataset (from E) with previously published proteome data from *in vivo* oxaliplatin-treated DRG (Yang et al., 2022). G. Rank-rank hypergeometric overlap between our DEP dataset (in E) and DEPs of the previously published oxaliplatin-treated proteome (Yang et al., 2022). DEP, differentially expressed protein; LFC, log_2_ fold change.

**Supplemental Figure 7.**
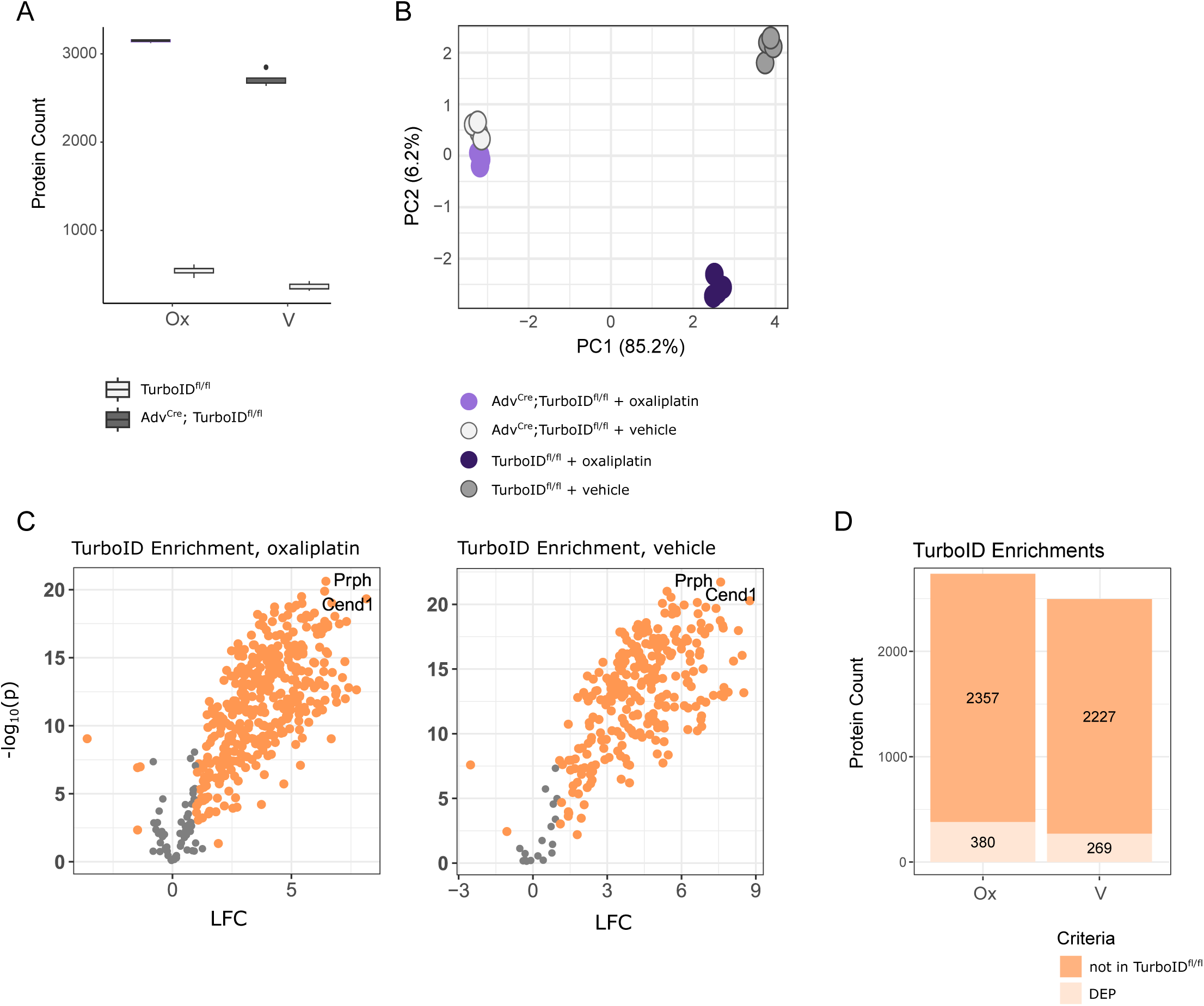
Proteomics of DRG explants from Adv^Cre^;TurboID^fl/fl^ and TurboID^fl/fl^ mice. A. Protein group counts across conditions. B. PCA across conditions. C. Differentially expressed proteins between Adv^Cre^;TurboID^fl/fl^ and TurboID^fl/fl^ explant samples, for both vehicle and oxaliplatin-treated explants. D. Numbers of enriched proteins, including both DEPs from C (Adv^Cre^;TurboID^fl/fl^ vs TurboID^fl/fl^), as well as proteins detected in Adv^Cre^;TurboID^fl/fl^ but not TurboID^fl/fl^ controls (“not in TurboID^fl/fl^”), please see methods for details. LFC = log_2_ fold change. Ox = oxaliplatin. V = vehicle.

## Supplemental Tables + Methods: Proximity labelling reveals the compartmental proteome of murine sensory neurons

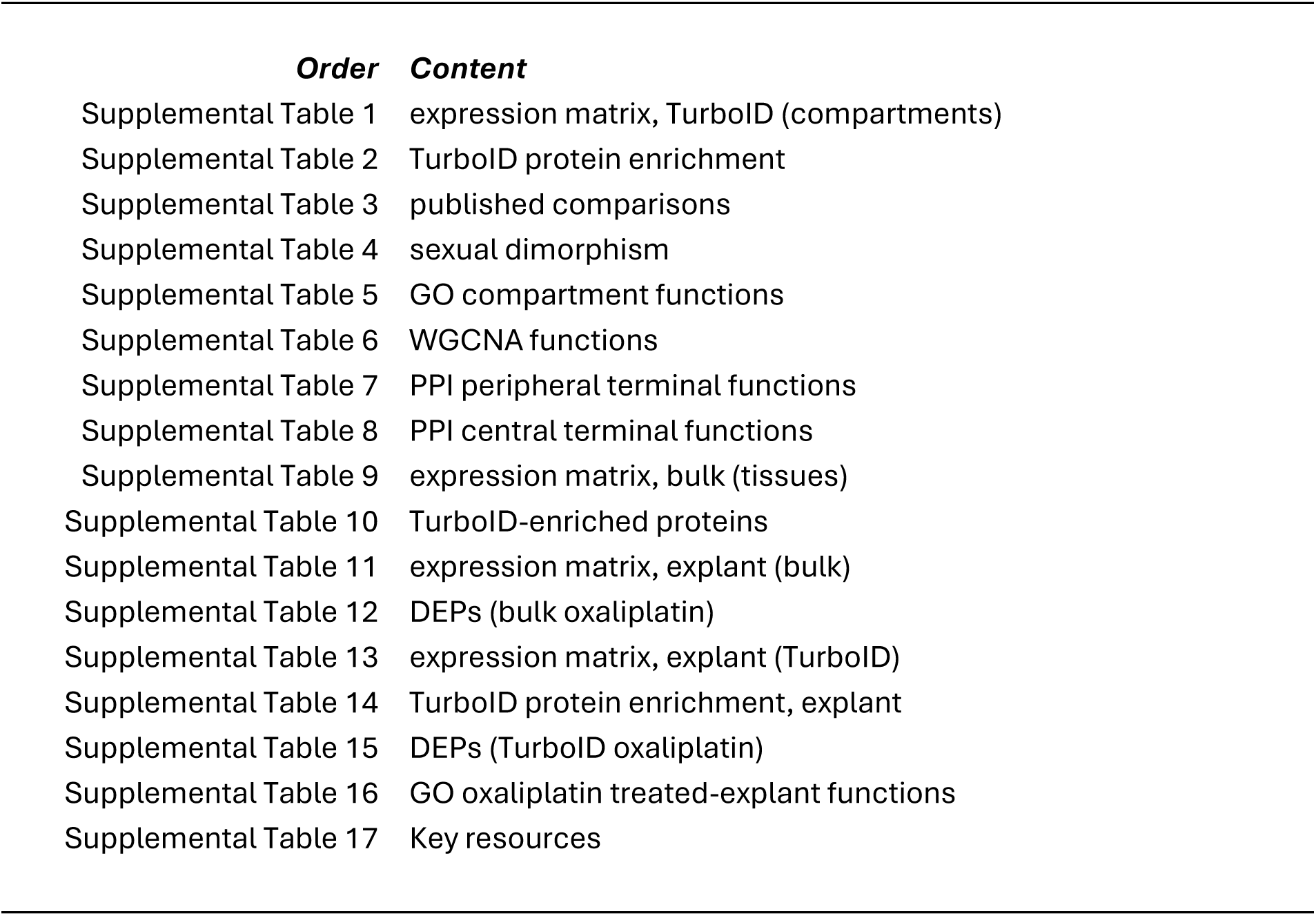

**Supplemental Table 17.**
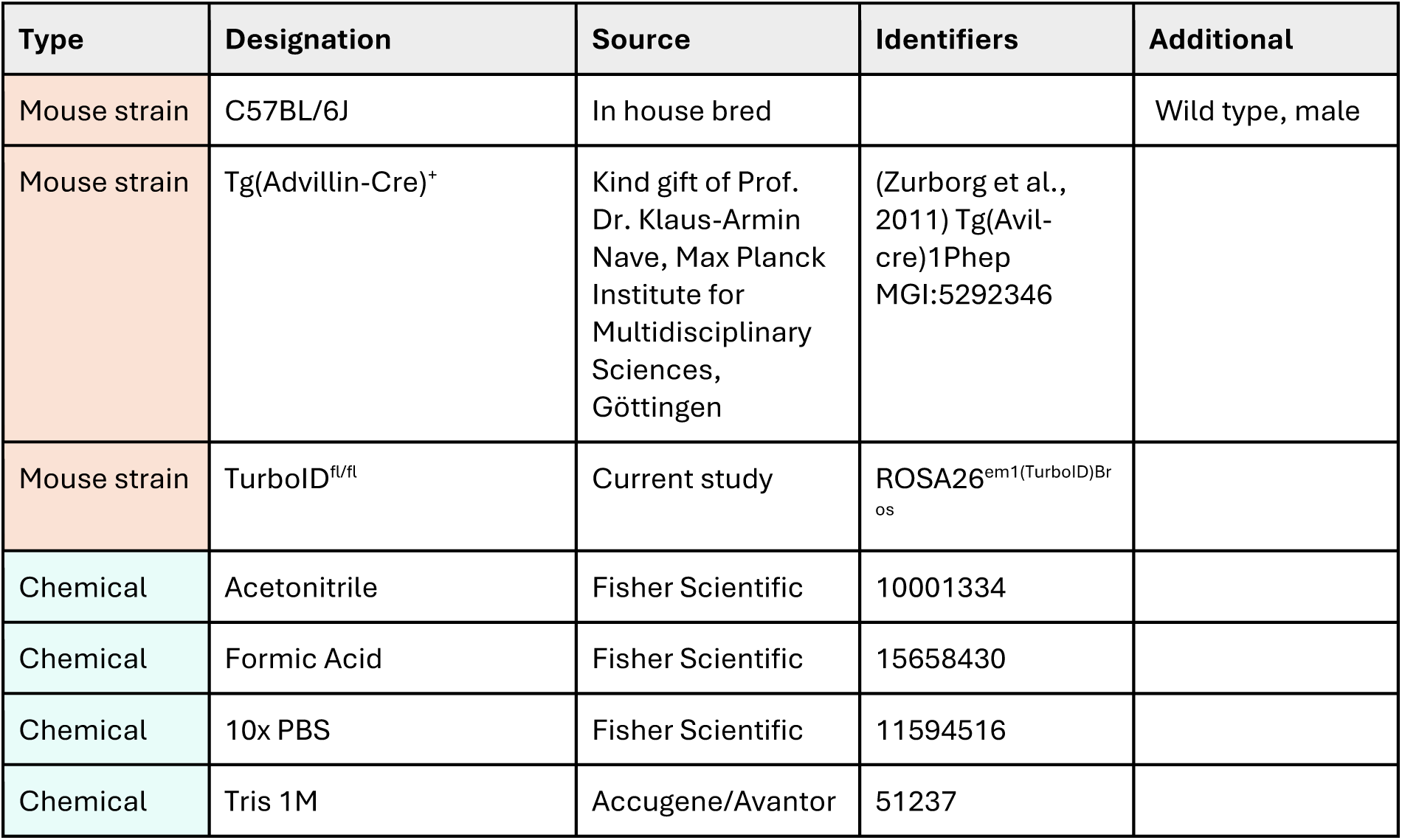
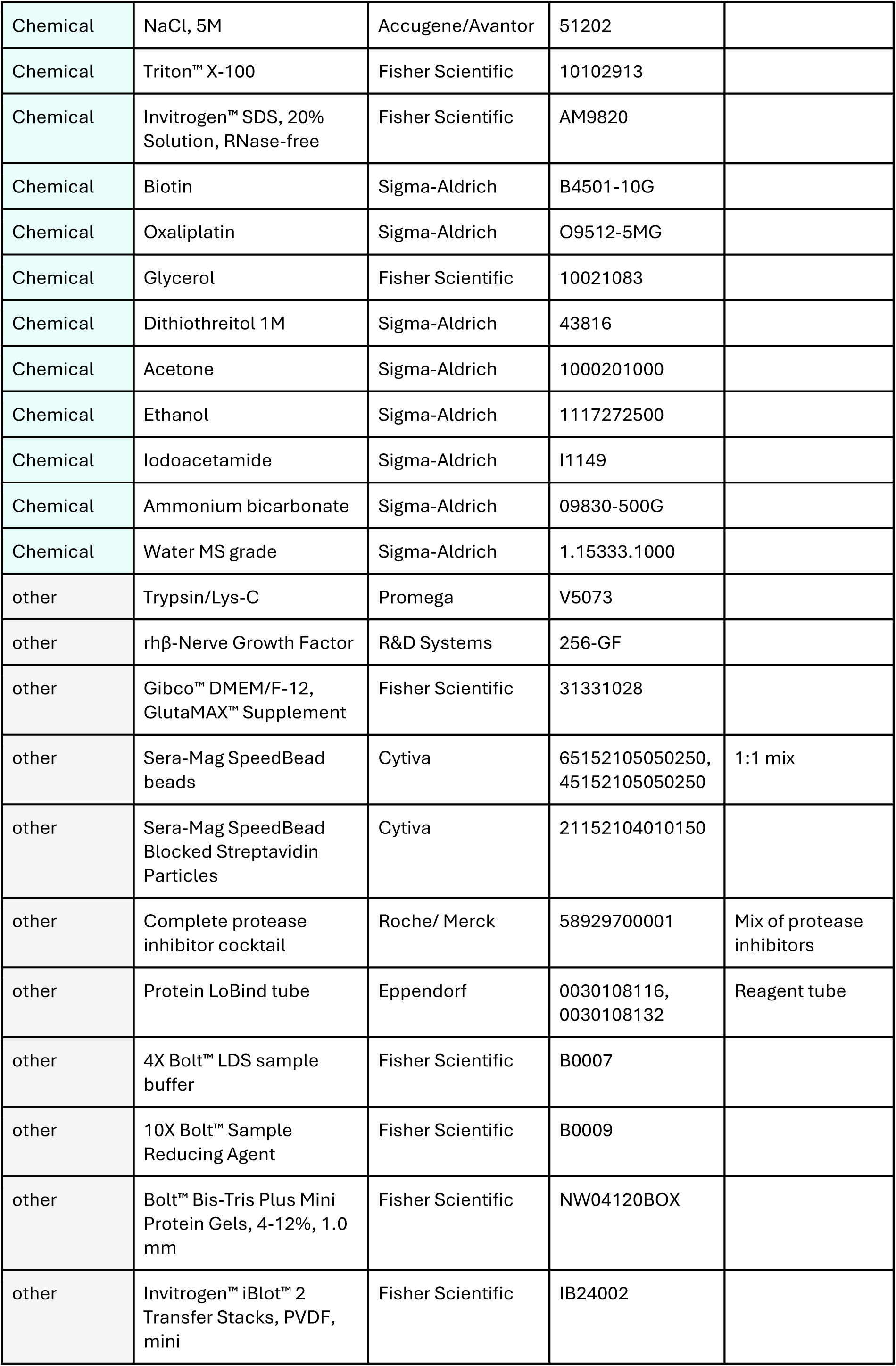
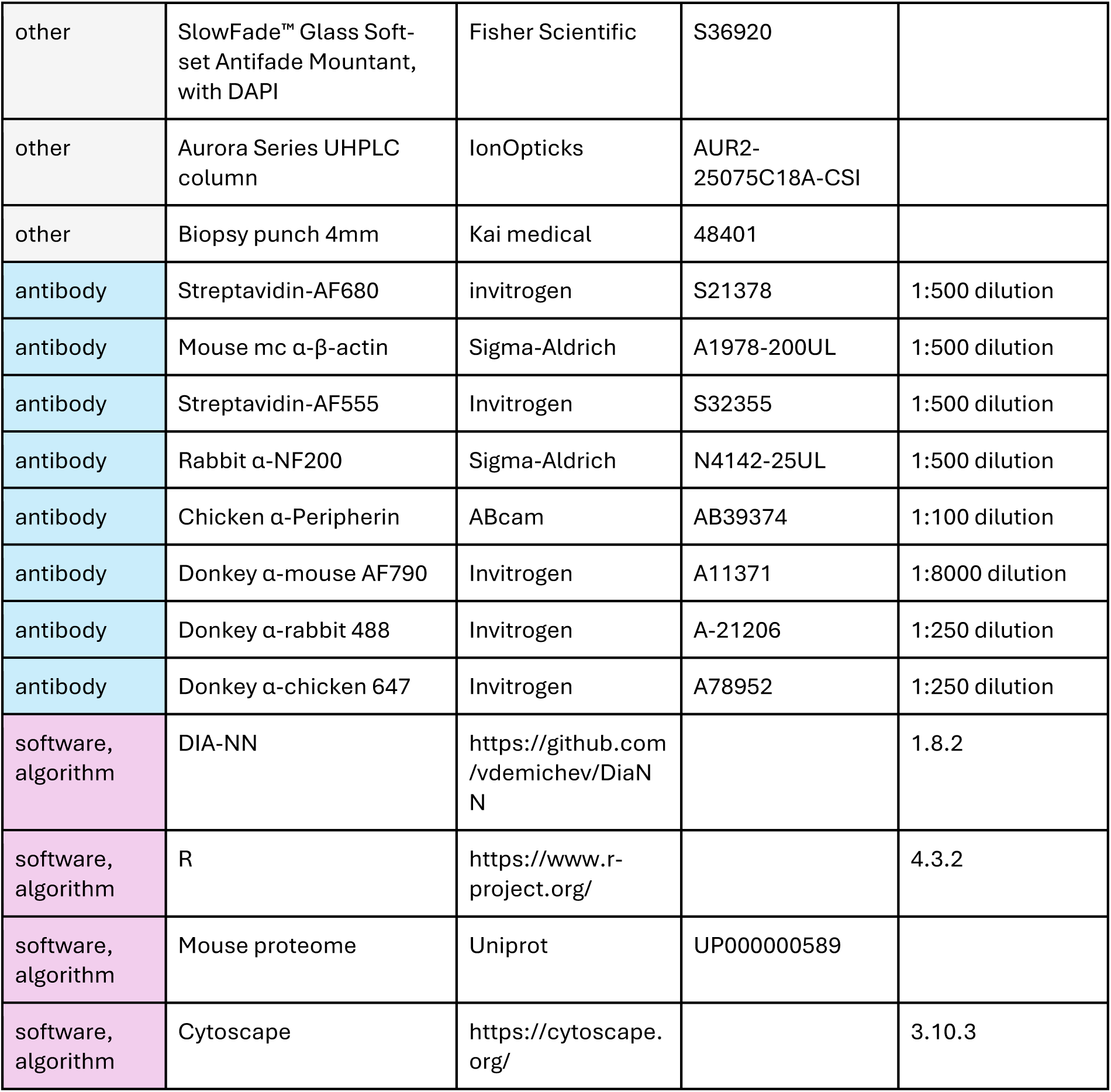
Key Resources.

## Supplemental Methods: Proximity labelling reveals the compartmental proteome of murine sensory neurons

### Generation of TurboID mutant mice using CRISPR/Cas9

The conditional TurboID knock-in (cKI) mouse line was generated by site-directed CRISPR/Cas9 mutagenesis in the ROSA26 locus. Superovulated C57Bl6/J females were mated with C57Bl6/J males and fertilized eggs collected. In-house produced CRISPR reagents (hCas9_mRNA, sgRNAs, preformed Cas9_sgRNA-RNP complexes and the homology-directed repair fragment (HDR fragment)) were microinjected into the pronucleus and the cytoplasm of pronuclear-stage zygotes using an Eppendorf Femtojet. Importantly, all nucleotide-based CRISPR/Cas9 reagents (sgRNAs and hCAS9_mRNA) were used as RNA molecules and were not plasmid-encoded, reducing the probability of off-target effects due to the short lifetime of RNA-based reagents (Tycko et al., 2019; Doench et al., 2016). The TurboID-containing HDR fragment was generated by subcloning a V5-tag-TurboID sequence into the Ai14-ROSA26-based plasmid backbone (Madisen et al., 2010) and isolated as a linear DNA fragment by SacI/AgeI restriction enzyme double digestion. Two sgRNAs targeting the ROSA26 Locus were selected using the guide RNA selection tool CRISPOR (Haeussler et al, 2016; Concordet & Haeussler, 2018). The correct locus-specific insertion of the HDR fragment was confirmed by localization PCRs with primers upstream and downstream of the HDR sequence.

Breeding Strategy: The Adv^Cre^;TurboID^fl/fl^ line was regularly maintained on a C57Bl/6 background. For all experiments, Adv^Cre^;TurboID^fl/fl^ males were crossed to TurboID^fl/fl^ females to generate Adv^Cre^;TurboID^fl/fl^ experimental mice (of both sexes) as well as Adv^Cre^ - negative TurboID^fl/fl^ littermate controls (TurboID^fl/fl^, of both sexes). We employed a four-tier strategy to prevent and assess potential germline recombination in line with published recommendations (Luo et al., 2020): (1) All breedings were set up with Adv^Cre^ present in male but not female breeders as suggested another published Adv^Cre^ knock-in mouse line (https://www.jax.org/strain/032536); (2) Adv^Cre^ was always maintained in a heterozygous state; (3) all mice were genotyped; (4) our genotyping protocol included a primer that would detect a potential Cre-recombined allele, which has not been the case in any progeny generated. Thus, to the best of our knowledge, we did not observe germline recombination in Adv^Cre^;TurboID^fl/fl^ mice.

### Genotyping Strategy

For TurboID genotyping, genomic DNA (gDNA) was isolated from tail biopsies using a genomic DNA isolation kit (Nexttec, #10.924). For the location PCRs, 20 μL reactions were prepared using 1 μL clean gDNA (15-80ng), 4 µL PrimerSet (4 pmol final each), 4 µL 5X Reaction Buffer (Finnzymes #F-524), 0.4 µL PhireHot-Start II Taq DNA Polymerase (Finnzymes #F-122L), 1 µL 10 mM dNTPs (Bioline #DM-515107), 4 µL Hi-Spec Additive (Bioline #HS-014101) and 5.6 µL H2O. Thermocycler parameters location-PCR Rosa26 left arm: 98°C for 5 min, (98 °C for 45 s, 64 °C for 30 s, 72 °C for 60 s) repeated for 34 cycles, final step at 72 °C for 10 min. Thermocycler parameters-location PCR Rosa26 right arm: 98°C for 5 min, (98 °C for 45 s, 68 °C for 30 s, 72 °C for 180 s) repeated for 34 cycles, final step at 72 °C for 10 min.

For the diagnostic routine genotyping PCR, 20 μL reactions were prepared using 1 μL clean gDNA (15-80ng), 4 µL PrimerSet (4 pmol final each), 4 µL 5X Reaction Buffer (Biozym #331620XL), 0.2 µL Hot-Start Taq DNA Polymerase (Biozym #331620XL), 1 µL 50 mM MgCl2 Sondermann, Barry et al. | 4 (AGCTLab stock) and 9.8 µL H2O. Thermocycler parameters: 96°C for 3 min, (94 °C for 30 s, 62 °C for 60 s, 72 °C for 60 s) repeated for 32 cycles, final step at 72 °C for 7 min.

### Rosa26 Locus sgRNAs

- Sense Rosa26 protospacer-sgRNA1 sequence: 5‘-ACTCCAGTCTTTCTAGAAGA-3‘ (PAM = TGG)
- Antisense Rosa26 protospacer-sgRNA2 sequence: 5‘-CGCCCATCTTCTAGAAAGAC-3‘ (PAM = TGG)

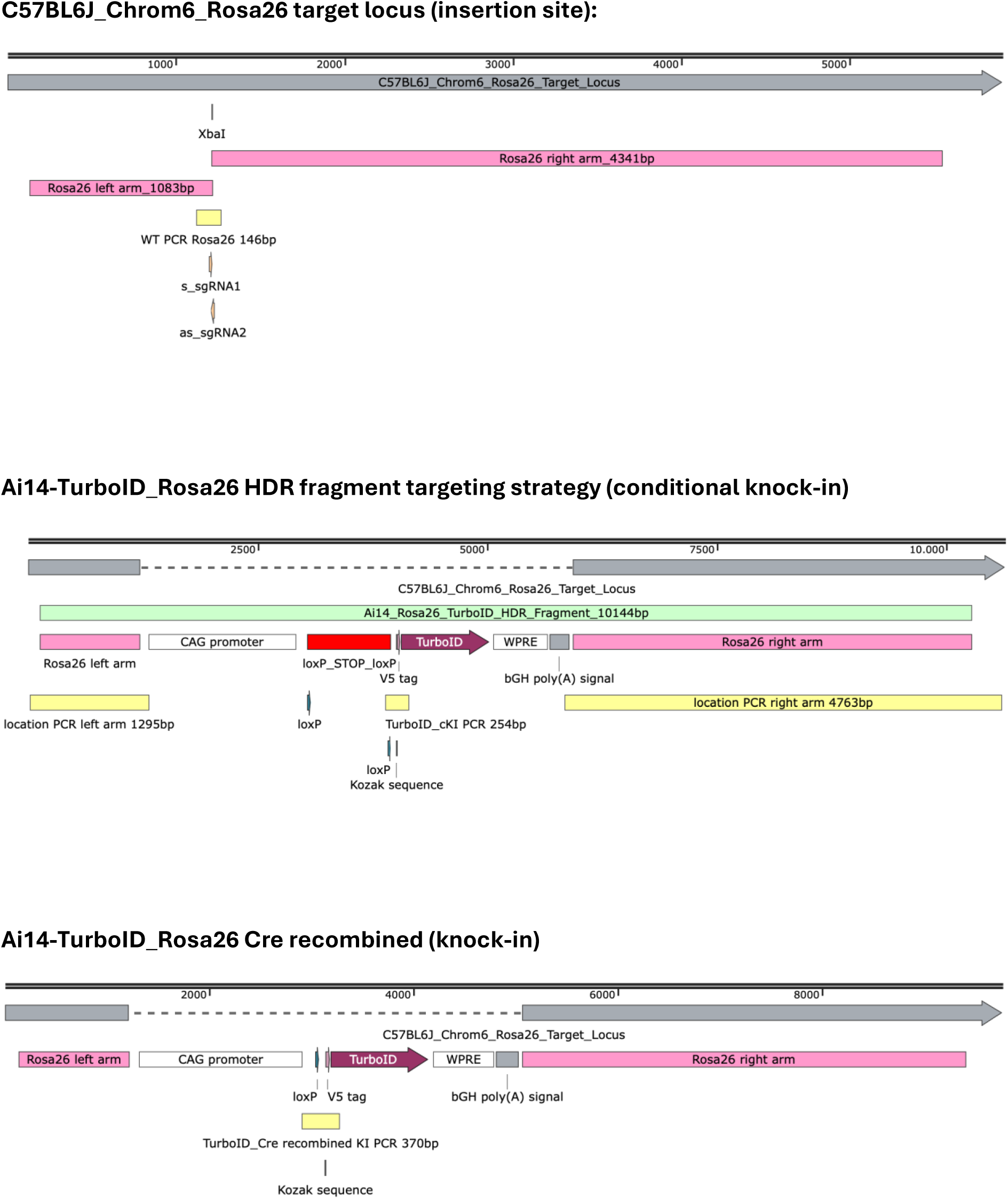

### SMethods, Figure1. Schematic representation of the target locus, the targeting strategy (HDR fragment) and the Cre recombination event

#### Ai14-TurboID_Rosa26 HDR fragment and sequence annotations

Lower case letters = upstream and downstream HDR fragment ROSA26 sequences Capital letters = HDR fragment after locus-specific insertion

Magenta = ROSA26 left- and right arm sequence Blue = CAG-promoter

**Red BOLD** = loxP sites

**<u>Underlined BOLD</u>** = loxP flanked STOP cassette **Green BOLD** = ATG Start- and TAA Stop codon **Brown BOLD** = V5-tag

**Purple BOLD** = TurboID

**Black BOLD** = WPRE

<u>Black underlined</u> = bGH poly(A) signal

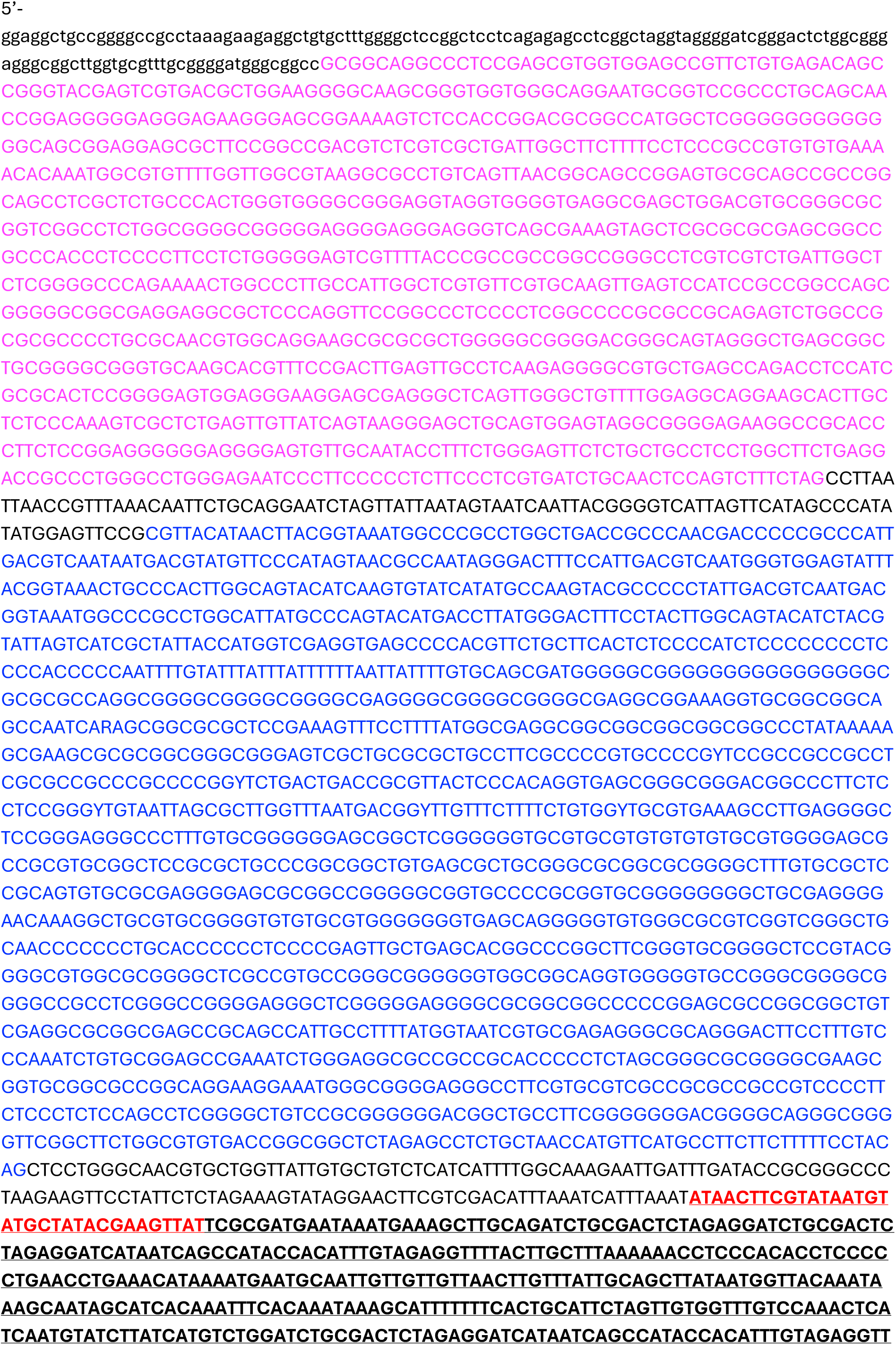
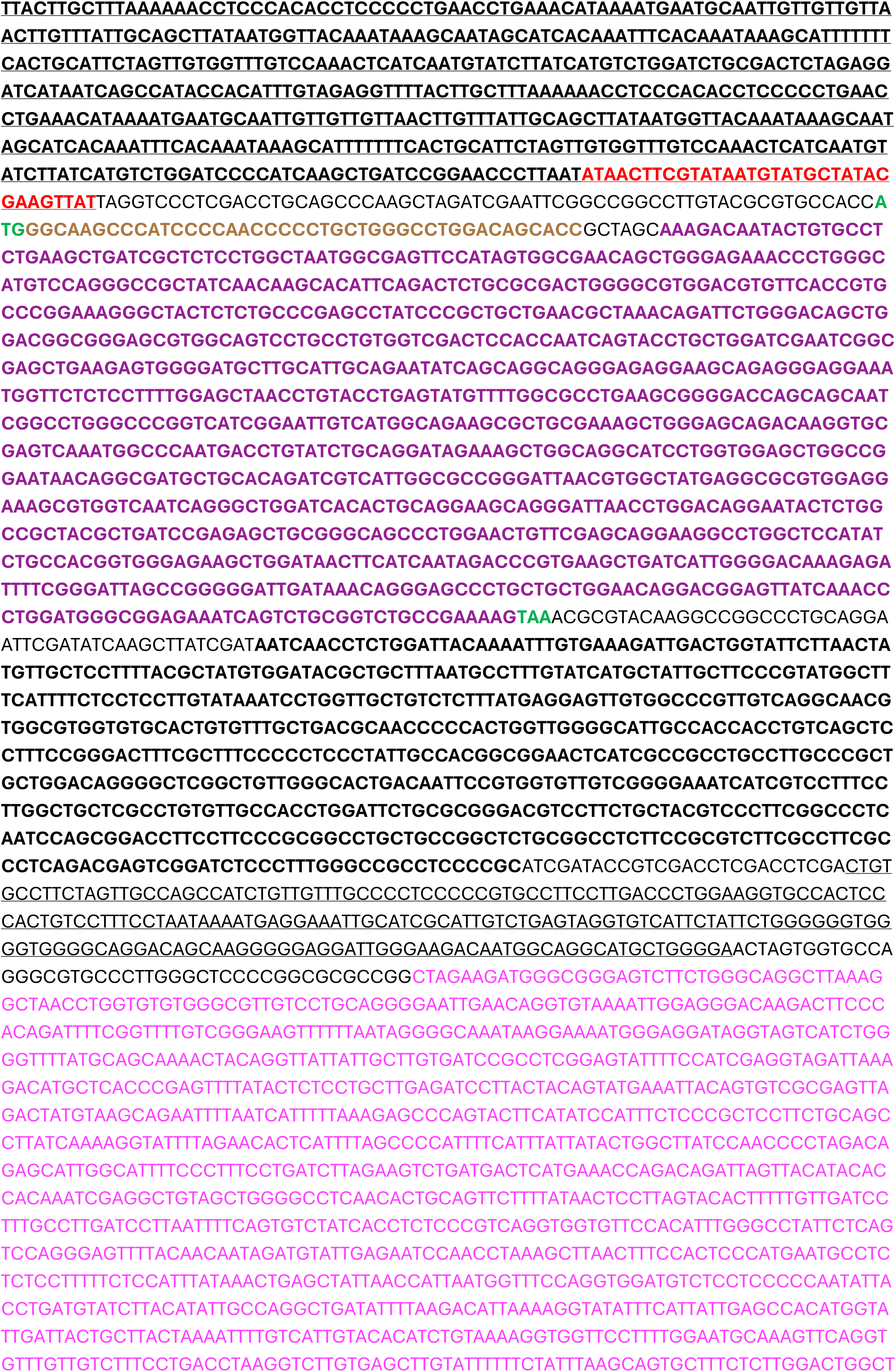
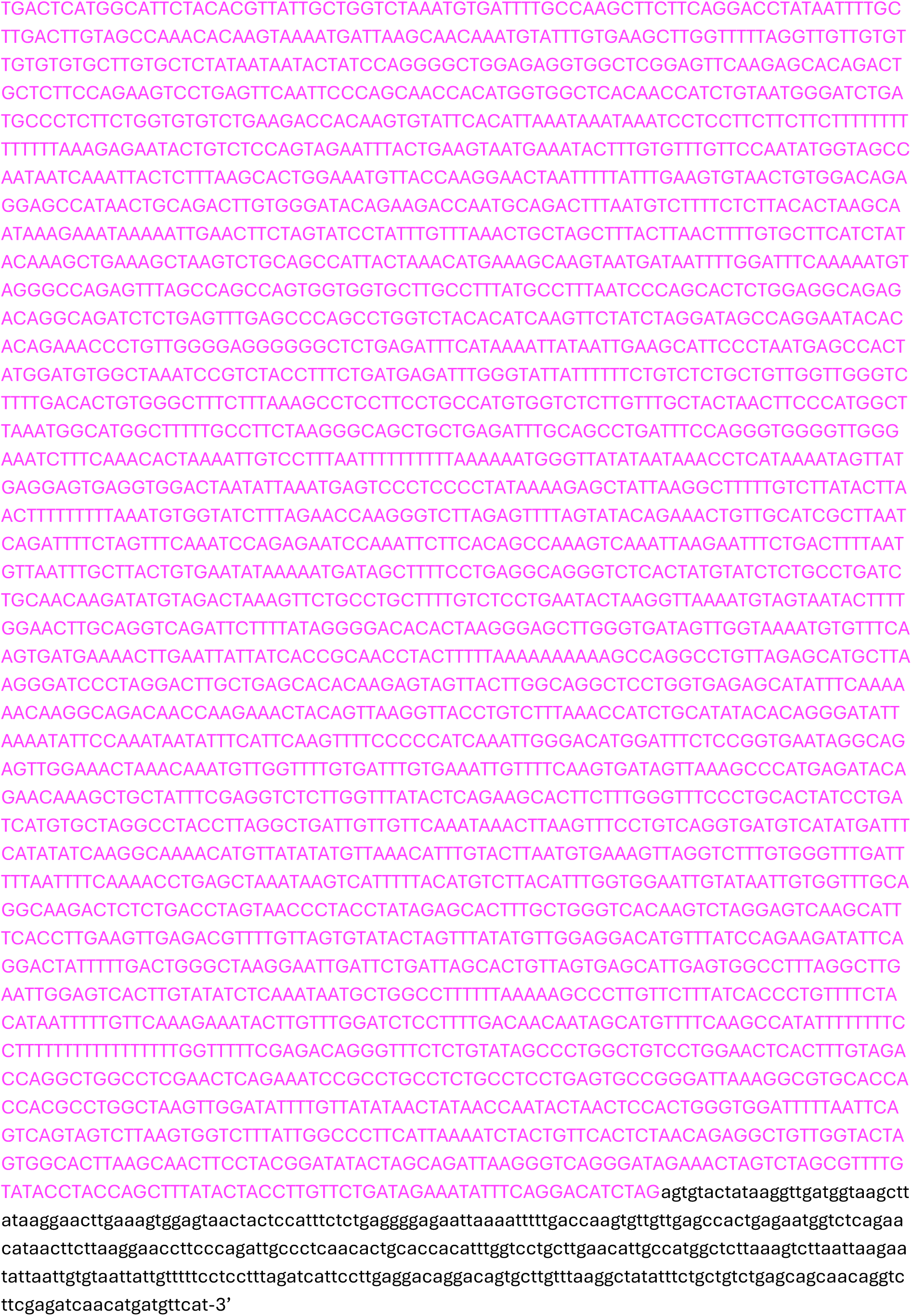

### **SMethods, Sequence.** Ai14-TurboID_Rosa26 HDR fragment and sequence annotations

Sondermann, Barry et al. | 9

### SMethods, Table. Primer sequence and fragment size information

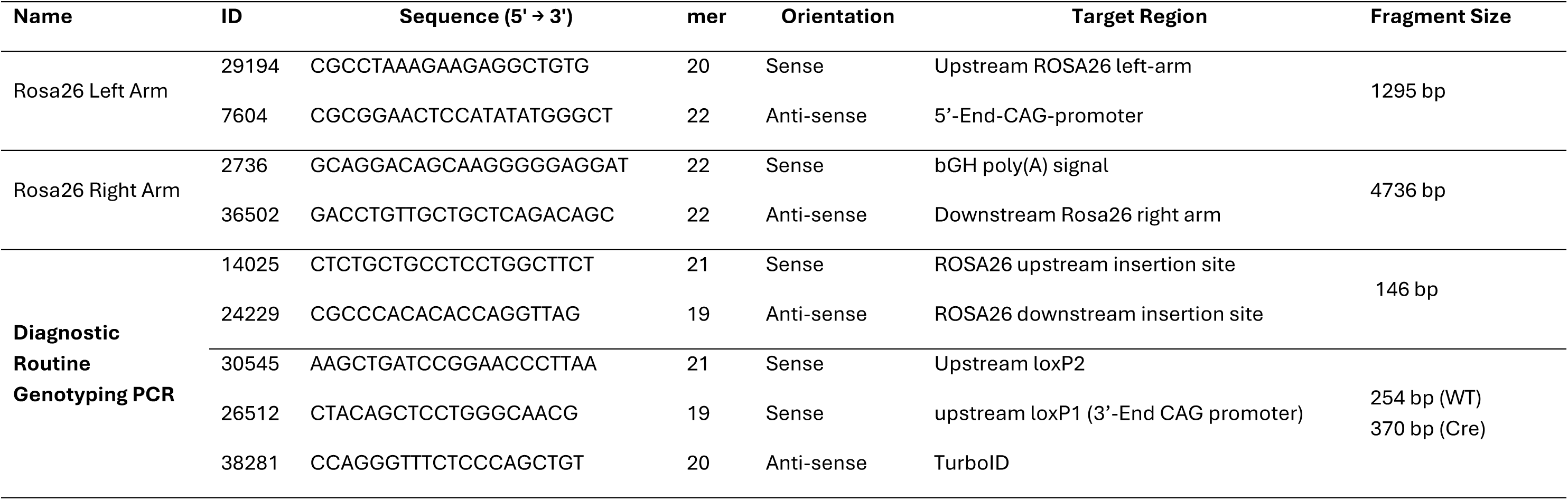

